# The Heart’s Pacemaker Mimics Brain Cytoarchitecture and Function: Autonomic innervation, a peripheral glial cell web, and a novel S100B expressing interstitial cell type impart structural and functional complexity to the sinoatrial node

**DOI:** 10.1101/2022.02.04.478900

**Authors:** Rostislav Bychkov, Magdalena Juhaszova, Miguel Calvo-Rubio, Lorenzo A. H. Donald, Chris Coletta, Chad Shumaker, Kayla Moorman, Syevda Tagirova Sirenko, Alex V. Maltsev, Steven J. Sollott, Edward G. Lakatta

## Abstract

**Objectives:** This study sought to describe the 3D cytoarchitecture of sinoatrial node tissue, including autonomic innervation, peripheral glial cells, and pacemaker cells.

**Background:** The sinoatrial node of the heart produces rhythmic action potentials (AP), generated via calcium signaling within and among pacemaker cells. Our previous work has described the SAN as composed of an HCN4-expressing pacemaker cell meshwork, which merges with a network of CX43+/F-actin+ cells. It is also known that sympathetic and parasympathetic innervation from epicardial ganglia create an autonomic plexus in the sinoatrial node, which modulates heart rate and rhythm. However, the anatomical details of the interaction of this plexus with the pacemaker cell meshwork have yet to be described.

**Methods:** 3D confocal laser-scanning microscopy of triple immunolabeled SAN whole mount preparations with combinations of antibodies for HCN4, S100B, GFAP, ChAT or VAChT, and TH, and transmission electron microscopy (TEM).

**Results:** The SAN exhibited heterogeneous autonomic innervation, which was accompanied by a web of peripheral glial cells (PGCs). Further, we identified a novel S100B+/GFAP- interstitial cell population, with unique morphology and distinct distribution pattern, creating complex interactions with other cell types in the node. TEM images showed a similar population of cells, here identified as telocytes, which appeared to secrete vesicles towards pacemaker cells. Application of S100B protein to SAN preparations induced distinct changes in rhythmogenic calcium signaling.

**Conclusions:** The autonomic plexus and its associated peripheral glial cell web, a novel network of S100B expressing interstitial cells resembling telocytes, and a meshwork of HCN4+ cells interact to impart structural complexity to the sinoatrial node.

**Summary Table:** 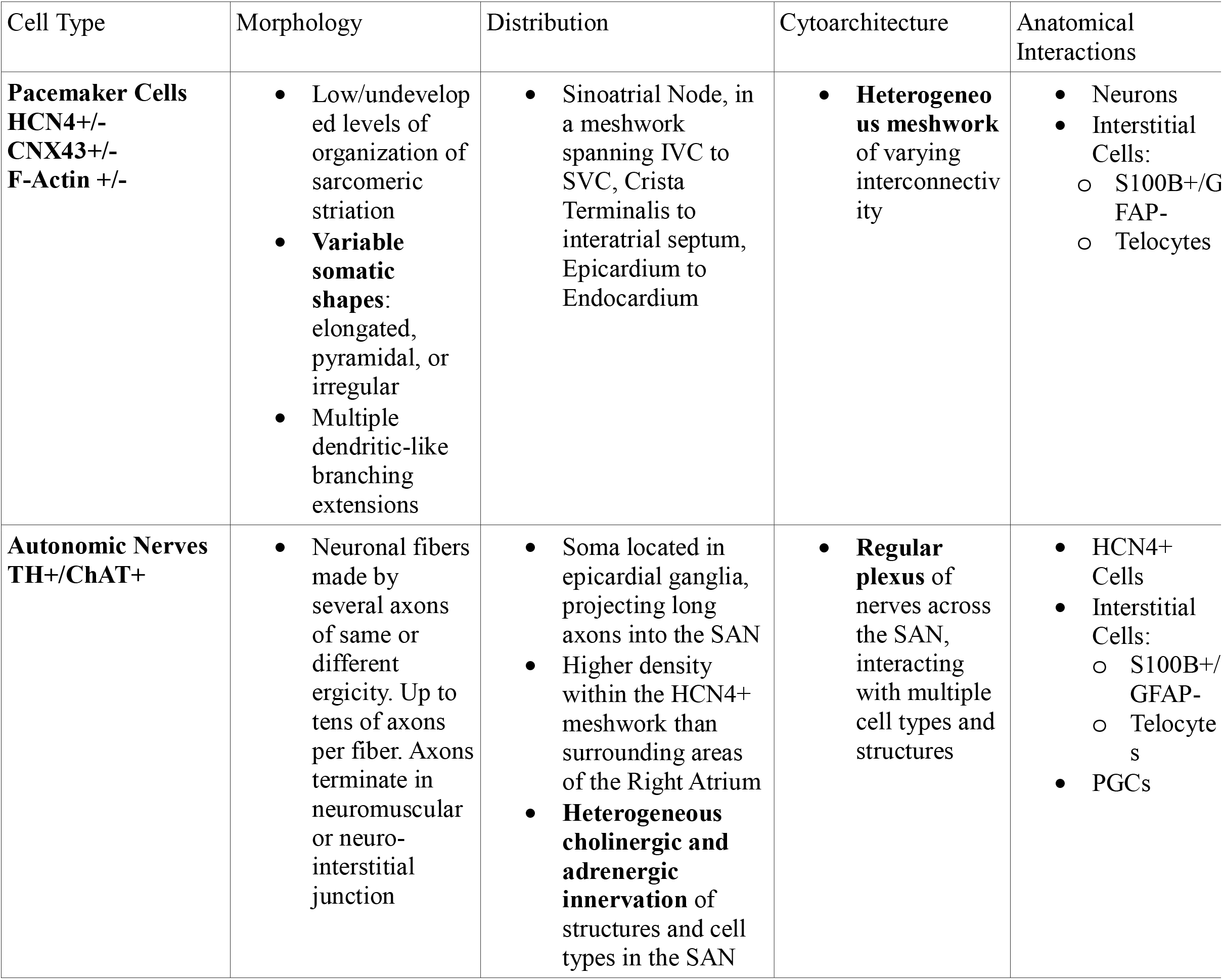

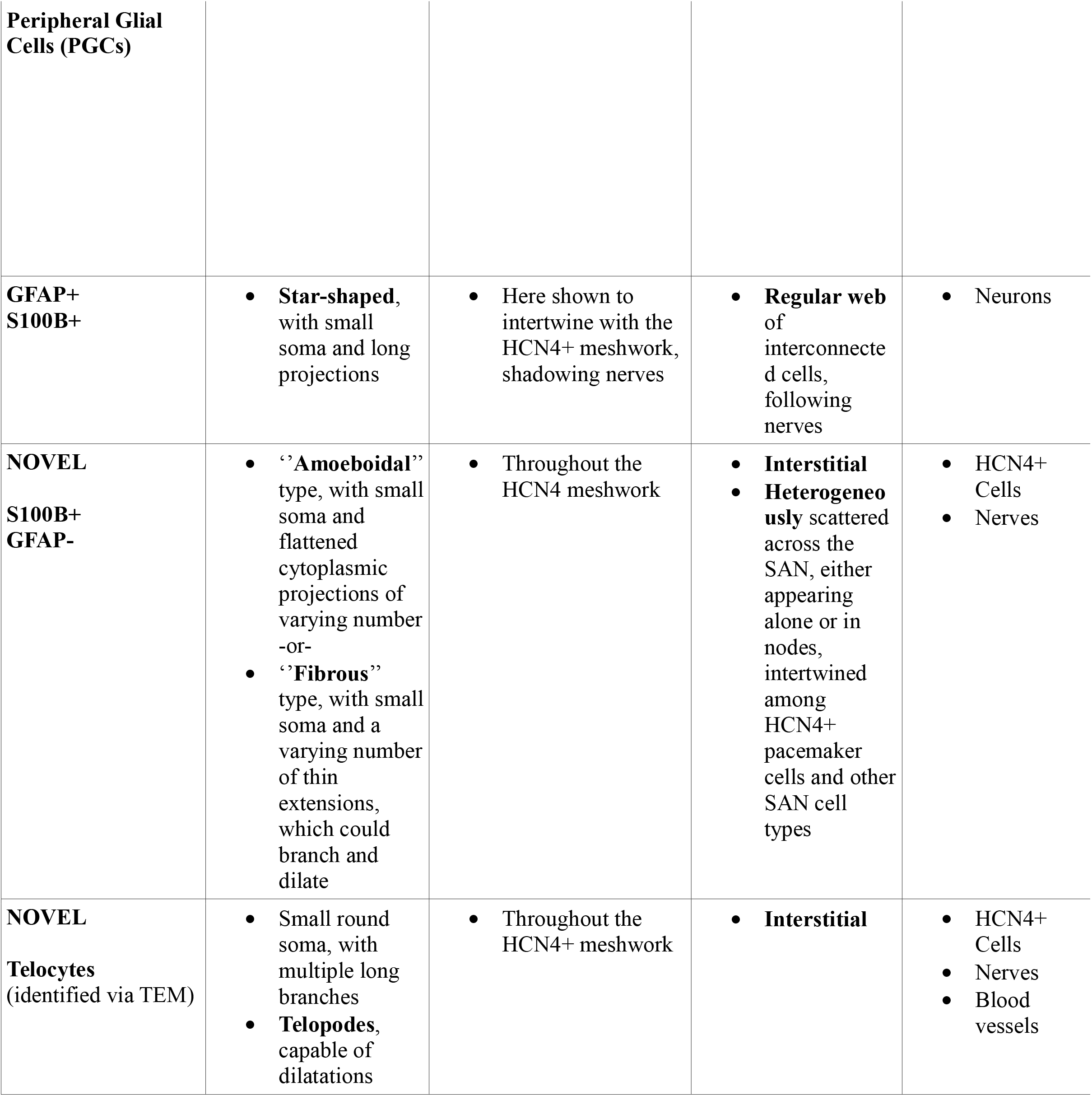

## Introduction

The heart is a central player in the hierarchical system of clocks operating within the body-wide neuro-visceral axis that determines the timing of synchronized rhythmic functions ranging from milliseconds to days, including cardiac beating rate. Brainstem neurons, with connections to and from cortical neurons, generate electrical signals that are conducted rostrally to neuronal ganglia embedded on the epicardial surface, and to blood vessels that emerge from the heart (1). Axons from epicardial ganglionic neurons, “the little brain on the heart” (2), penetrate into the sinoatrial node (SAN; (3)), encountering and embracing SAN pacemaker cells, secreting neurotransmitters. Activation of autonomic receptor driven signaling within SAN cells by these transmitters impacts the AP firing rate and rhythm in SAN cells by modulating the tempo and ticking speed of an intrinsic coupled-clock system that drives their automaticity: the sarcoplasmic reticulum, a Ca^2+^ clock that generates diastolic local sub-cellular Ca^2+^ signals that self-organize via a criticality mechanism (4–6) and couples to an ensemble of surface membrane ion channels (M-clock), operating on a limit cycle mechanism to produce plasma membrane current oscillations (7, 8). Evolution of the resultant electrochemical signal produced by the coupled-clock system during diastole results in progressive depolarization of the cell surface, culminating in an action potential. Neurotransmitter input to SAN cells impacts on the degree to which the Ca^2+^ and membrane potential clocks couple to each other, and therefore the duration and frequency of the AP ignition process, by modulating the kinetics of the molecular functions that drive the coupled clock system in the absence of neurotransmitter input. For example, sympathetic autonomic receptor stimulation improves clock coupling, resulting in an earlier onset of the ignition process and an increase in the pacemaker AP firing rate, in contrast to parasympathetic stimulation.

A recent discovery has added an entire new layer of complexity to the initiation of each heartbeat that extends well beyond the coupled-clock system intrinsic to individual SAN cells. Specifically, local oscillatory calcium signals that are heterogenous in phase, amplitude and frequency emerge within a meshwork of hyperpolarization activated cyclic nucleotide gated potassium channel 4 (HCN4) expressing cells extending nearly the entire length and depth of the central SAN (9). Local Ca^2+^ signals that emerge within and among SAN pacemaker cells self-organize to create impulses that exit the SAN and are conducted to other parts of the heart to generate heart beats, even in the absence of input from the brain stem.

But, how local Ca^2+^ signals within and among SAN pacemaker cells can self-organize into a critical synchronized event that emerges from the SAN to initiate heartbeats in the absence of neuronal input is not at all intuitive. Recent observations provide a valuable clue: it has been noted that this multi-scale, complex process of impulse generation by pacemaker cells residing within the SAN resembles the emergence of organized signals from heterogeneous local signals generated occurring within clusters of neurons comprising brain neuronal networks. The variable morphology of HCN4+ pacemaker cells within the SAN suggests that these cells may differ in the fine tuning of their functions that could contribute to heart rhythm generation in different ways. Furthermore, some HCN4^+^ cells project branches to the neighboring cells ending with end feet (9) that are similar in shape and size to those of astrocytes within the brain (10). Because the communication between SAN cells (9) resembles the emergence of neuronal signals within the brain (11), we reasoned that, beyond the known association of PGCs and autonomic nerves, the SAN also harbors multicellular complexes similar to the neuro-glial interactions in nervous tissue.

Brain glial cells are active players in the formation and function of neuronal brain circuitry, and the incompleteness of discussing brain structure and function without the inclusion of glial cells has been realized previously (12). The role of glial cell support functions including myelination, synapse-pruning, and macrophagy has long been known, and their crucial function to modulate neuronal communication has also been appreciated (13). For instance: microglia have been shown to reduce neural activity by releasing adenosine, in a mechanism similar to inhibitory synaptic transmission (14); and Satellite glia were shown to regulate the strength and firing rate of spontaneous synaptic transmission between sympathetic neurons and to promote the formation of synaptic sites with a specific effect on the formation of presynaptic structures (14, 15). In contrast to the mass of information available on the roles of CNS glia, information regarding the presence of cytoarchitecture and functions of peripheral glial cells (PGC) in the heart is conspicuously sparse (16).

In our prior study we had discovered a novel, microscopic Ca^2+^ signaling paradigm of SAN operation: synchronized APs emerge from heterogeneous subcellular subthreshold Ca^2+^ signals, that resemble the multiscale complex processes of impulse generation within clusters of neurons in neuronal networks. Here, we hypothesized that neuronal cells, peripheral glial cells and pacemaker cells may interconnect in functional units within the SAN, and display a type of cytoarchitecture with anatomical interactions similar to those found in neuronal tissue. Specifically, glial and neuronal cells may create a microenvironment around pacemaker cells dedicated to the regulation not only of LCRs within individual pacemaker cells but also to the regulation of intercellular communication among pacemaker cells that crucially impact heart rhythm. The spatial pattern of LCRs across SAN that emerges from intercellular communications may be monitored and regulated by a functional unit of neuronal, glial and pacemaker cells. To this end, we utilized a novel 3D confocal tile imaging techniques to optically dissect the entire mouse SAN (top to bottom, inside to outside) and to reconstruct SAN cytoarchitecture from 3D optical slices, with respect to neuronal fibers and peripheral glial cells.

## Methods

Detailed methods for immunolabeling, confocal microscopy, TEM, and data analysis are described in the **Supplemental Information**.

### Antibodies

HCN4^+^ cells were identified by rabbit polyclonal antibodies for hyperpolarization activated, cyclic nucleotide-gate cation channels HCN4 (1:300, Alomone Labs, Jerusalem, Israel). S100B meshwork was identified by a recombinant rabbit monoclonal anti-S100B antibody (1:250; clone EP1576Y; catalog # ab52642; Abcam, Cambridge, MA), and a chicken polyclonal anti-S100B antibody (1:300; catalog # 287006; Synaptic Systems, GmbH, Göttingen Germany).

The parasympathetic nervous system was visualized by a goat polyclonal anti-choline acetyltransferase antibody (ChAT; 1:300; cat. # AB144P from MilliporeSigma, Burlington, MA) and a guinea pig anti-vesicular acetylcholine transporter (VAChT 1:300; cat # 139105 from Synaptic Systems). Sympathetic nerve fibers were visualized by two anti-tyrosine hydroxylase (TH) antibodies, a mouse monoclonal (clone LNC1, Alexa-488 conjugated, cat # MAB318-AF-488, 1:250 dilution), and a chicken polyclonal (AB9702, 1:250), both from MilliporeSigma. Astrocytic glial cells were labelled with two polyclonal anti-GFAP antibodies; a goat anti-GFAP (1:300, cat # SAB2500462 from MilliporeSigma), and chicken anti-GFAP (1:300; cat # PA1-10004, Invitrogen) and a rabbit anti-S100B (1:300; cat# ab196175 from Abcam). Nuclei were visualized with 500 nM DAPI (4′,6-diamidino-2-phenylindole) in PBS for 30 min.

### Imaging of local Ca^2+^ signals in SAN tissue

These experiments were performed in ex-vivo, horizontally-mounted, perfused SAN prep that was carefully dissected from mouse hearts. We used a 5x air objective to obtain a wide-field view of the SAN, and recorded Ca^2+^ dynamics using HCN4-GCaMP8 transgenic mice expressing a genetically encoded fluorescent Ca^2+^ indicator in HCN4+ pacemaker cells (The Jackson Laboratory; Strain #028344; for specific details about resolution, see **Supplemental Methods**). Background Ca^2+^ fluorescence was negligible in the diastolic phase, creating high signal-to-noise ratio when recording action potential-induced Ca^2+^ transients (APCTs) (**Fig. 15**). Our 2D-calcium images were recorded from the endocardial side of the SAN, and did not include Ca^2+^ dynamics within deeper layers of the HCN4+ meshwork or within pacemaker cells located near the epicardial surface. The signal-to-noise ratio of Ca^2+^ induced fluorescence allowed us to reliably detect APCTs from SAN preparations for up to 1 hour after the first moment of exposure to the excitation light of the objective during recording (n=3). In each preparation, S100B was added to the superfusate within 10-15 minutes from initial control recordings, and its effects were imaged.

## Results

### Panoramic imaging of the HCN4+ meshwork and intertwining neuronal network within the SAN

We optically sectioned SAN tissue using a conventional confocal microscopy technique and tiled 3D Z-stacks into a panoramic 3D image of the SAN 350 µm deep, 8 mm long and 4 mm wide images (**Fig. 1**) (for more detailed description of the confocal methodology and 3D reconstruction, see **Supplemental Methods**). The orientation of the semi-intact preparations (SAN + atria) was identical that in (9). We used the previously proposed nomenclature of “head”, “body” and “tail” (17) throughout the text to describe HCN4-immunoreactive (**red**) pacemaker meshwork. The “head” of the meshwork forms around the root of SVC, and continues through the “body” of the meshwork along the course of SA nodal artery, with the “tail” extending to the root of IVC. The panoramic 3D-image depicted in **Fig. 1** highlights innervation of the HCN4+ pacemaker meshwork interwoven with cholinergic and adrenergic fibers that form 3D SAN neuronal plexus. Pacemaker cells, parasympathetic fibers, and sympathetic fibers of the SAN neuronal plexus immunoreactive for HCN4, VChAT, and TH, respectively, were readily identifiable up to 350 µm in depth in the SAN tissue in all whole mount preparations (**n=3**). Compared with atrial areas adjacent to the root of SVC, the density of nerve fibers within the HCN4-positive meshwork of pacemaker cells was 2–3 fold higher (**n=3**) consistent with previous publications (18, 19).

**Fig. 1.**
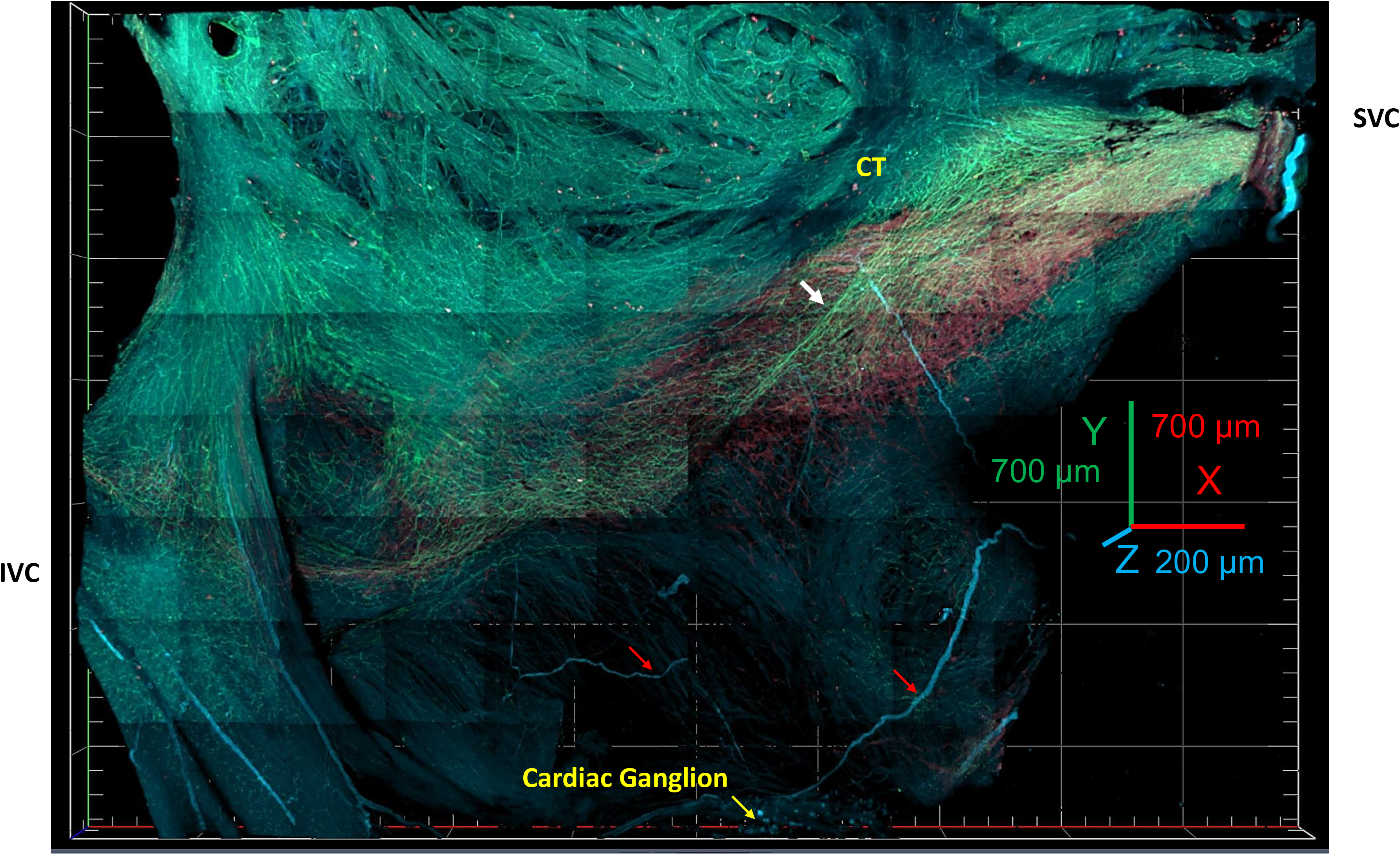
3D-image tiled from series of Z-stacks reconstructed from 2D-images obtained by confocal microscope optical slicing of a whole-mount SAN preparation. The image shows the area from the superior vena cava (SVC right side of the image) to the inferior vena cava (IVC left side of the image) and from the septum to the crista terminalis. The “head” of the HCN4-positive pacemaker cell meshwork forms around the root of the SVC; the “body” of the meshwork continues along the course of SA nodal artery, forming the “tail” that extends to the root of IVC. Most of the ventricles was dissected in order to flatten the atria and the SAN for imaging. The roots of the pulmonary vein remained intact but were outside of the image. The image presents a complete picture of all nerve fibers within each point of both the SAN and right auricle. The plexus of VAChT+ cholinergic (green color) and TH+ adrenergic (cyan color) innervation enwrapped the HCN4^+^ immunoreactive meshwork (red color) in both the SAN and pectinate muscles of the right auricle. Cardiac ganglia (yellow arrow at bottom of the figure) were located near the epicardial side of the septum. Red arrows point to nerves emanating from these ganglia, which penetrated the tissue. A cholinergic bundle of high-density neuronal fibers crossed in diagonal the body of the SAN from upper right to the lower left part.

Thus, 3D SAN panoramic imaging presents a near complete picture of the HCN4+ meshwork and all nerve fibers throughout both the SAN and right atrium. The general pattern and gradients of innervation in these panoramic images were used for orientation with respect to selecting regions of interest for detailing the fine structure of the HCN4+ meshwork and interwoven neuronal plexus at higher resolution.

### Intrinsic Cardiac Ganglia

Although our main focus was to characterize glial cells within SAN tissue, we began by examining the intrinsic cardiac ganglia (2), and following of the nerves that emerge from them and penetrate the SAN. Intrinsic cardiac ganglia (ICG) with VAChT- and/or TH-immunoreactive neuronal somata were located on the epicardial surface, close to the septum. We did not examine all ICG in our SAN preparations; ganglia served as a positive control for VAChT and TH immunolabelling. Intensity of immunolabeling together with the outline of the neuronal somata and processes indicated successful immunolabelling of the neuronal plexus within the SAN (**Fig. 2**). Closer inspection showed that diameters of the examined ICG in our preparations varied from 40µm to 150µm, consistent with previous reports (17,18,20). On average, mouse cardiac ganglia contained about 106±29 neuronal somata; while both VAChT- and TH+ neuronal fibers were detected within any given ganglionic plexus, cholinergic innervation was dominant. VAChT immunoreactivity was characteristic of the majority (**about 60%-80%**) of the neurons identified within ICGs (**Fig. 2 Panel A**), which is also in accordance with previously published data (20).

**Fig. 2.**
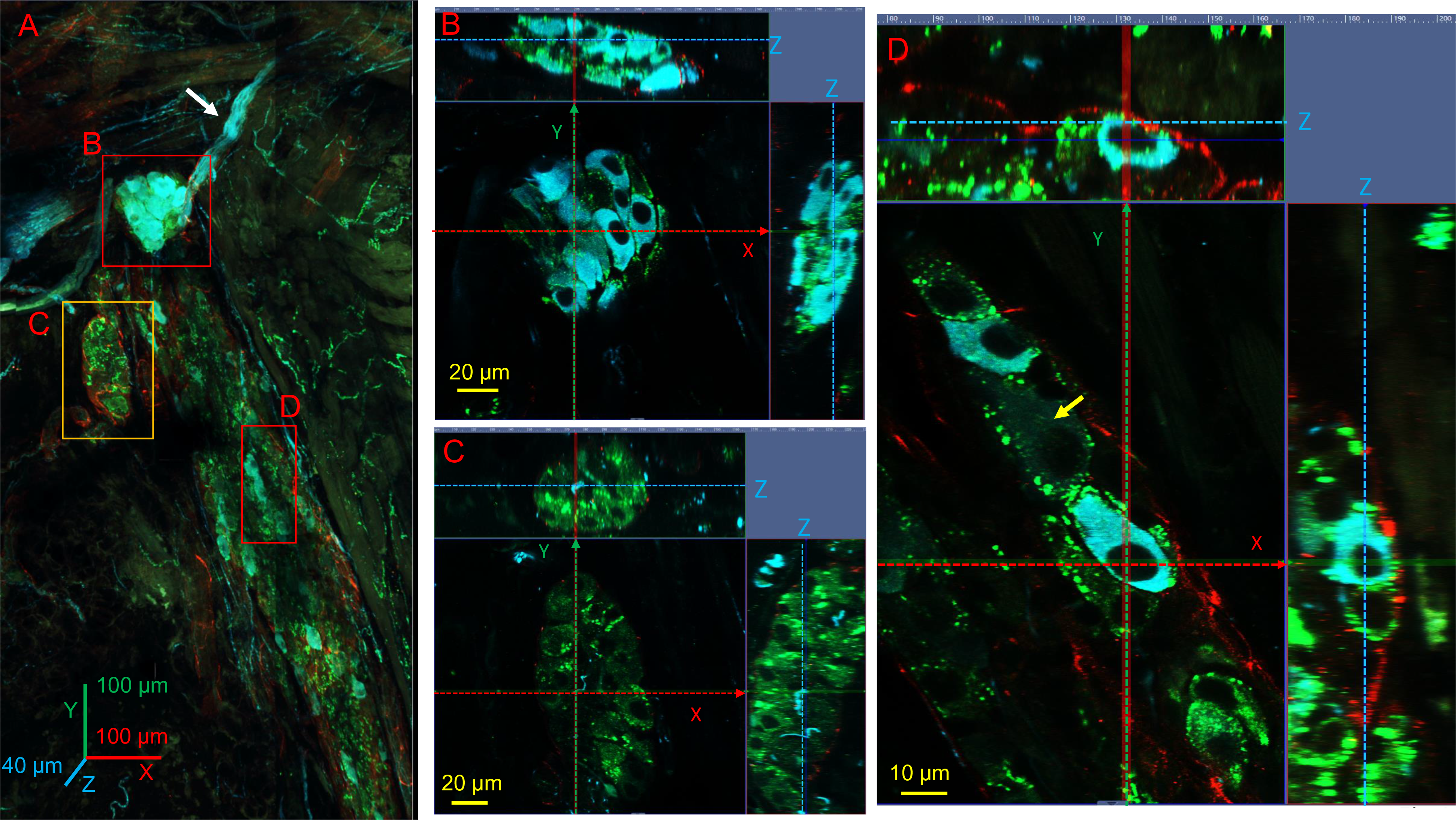
3D-image of intrinsic cardia ganglia tiled from a series of Z-stacks of the whole-mount SAN preparation. **Panel A** shows a series of 2D-images, obtained by confocal optical slices, that were used to reconstruct a 3D-image of the Z-stacks. The 3D-image in **Panel A** illustrates a group of three ganglia, two small ones (**Frames 2, 3**) and a portion of a larger interconnected one (**Frame 4**). Other ganglionic nerves are outside the field of view of the 3D-image. Vesicles immunoreactive to VAChT (**green color**) can be seen within the neuronal somata within all ganglia. One of the smaller ganglia, about 100μm in radius, with intracardiac nerve (**yellow arrow**) exiting the ganglion on the epicardial surface and going to the interior SAN tissue, contained primarily TH+ nerve fibers (**cyan neurons within frame 2**). The second smaller ganglion (**within frame 3**), about 50μm wide and 100μm long, contained predominantly VAChT immunoreactive neurons. The large elongated ganglion, about 100μm wide and 750μm long, lying adjacent to the ganglia in **frames 1 and 2**, surprisingly contained colocalized cholinergic and adrenergic neurons, as well neurons belonging to neither phenotype. The framed regions of interest in **Panel A** are shown with higher optical zoom in **Panels B, C** and **D** respectively. **Panels B, C** and **D** include: **a)** one 2D-image in the center of the panel that shows an example of a single optical slice through the Z-axis of the SAN, and **b)** two additional images, one on the top and another on the right side of the 2D-image, which illustrate the virtual Z-cut from the top to the bottom of the ganglion at the given X and Y position. The position of the optical cut of the Z axes is shown by red and yellow dotted lines in the side images. The X and Y positions taken for the virtual Z-cut are shown as blue dotted X and Y axes plotted within the central image. VAChT and TH immunoreactive varicosities are readily detectable within the ganglionic neuropil. Note that some neurons within ganglia in **Panel D** illustrates a ganglion with both VAChT and TH immunoreactive neurons as well as neurons immunonegative to VAChT and TH (**yellow arrow**).

Optical slicing of the IC ganglia showed that both TH and VAChT neurons were surrounded by varicose neural terminals which expressed VAChT (**Fig 2. Panels B, C, D**). Immunostaining of neural varicosities within the ganglionic plexus was more intense than the cytoplasmic and perinuclear regions of cholinergic ganglionic cells. Some neurons exhibited exclusively TH immunoreactivity, which varied among individual neurons and was not dependent on the neuron’s particular location inside the ganglion. VAChT and TH positive neuronal somata varied in size and shape. The purely adrenergic neurons were larger than their cholinergic counterparts, the long axis of VAChT neurons averaged 19.6±0,8μm in average, while that of the TH-neurons averaged 22.5±1.2μm. We also identified neurons within IC ganglia that were neither VAChT or TH positive, suggesting the presence of other neurotransmitters within ganglionic neurons, in accordance with previous publications (20). Interestingly, an HCN4+ membrane surrounded the ICG (**Figure 2. Panel D)**.

### 3D density gradients in cholinergic and adrenergic innervation of the pacemaker meshwork

3D-images were used to visual isotropic or anisotropic 3D-density pattern of the neural plexus in relation with the HCN4+ meshwork of pacemaker cells, as determined by the number of DAPI-stained nuclei of immunolabeled autonomic cells within a given area of tissue. The meshwork shown covers area about 7mm long, 4mm wide, and 300µm in depth. The panoramic 3D-image shown in **Figure 3** covers a SAN area of 2mm by 2mm. It was tiled from Z-stacks of 2D images (300µm by 300µm) that have been recorded with confocal microscopy by optical slicing of the whole-mount heart preparations. These comprehensive images were taken in the “body” of the SAN close to the SAN artery and show all immunoreactive cells and fibers in the tissue. Elaborate details of the neural plexus captured by the 2D scans were used to construct 3D-images from the Z-stacks, creating a high-resolution 2mm by 2mm panoramic image. The 3D-images in **Fig. 3** visualize the three-dimensional cytoarchitecture of the HCN4+ pacemaker meshwork and neural plexus which can be described in terms of 1) cellular shapes and densities; 2) orientation of the pacemaker cells and neuronal fibers with respect to epicardium, endocardium, SVC, and IVC; 3) apparent layering of pacemaker cells and HCN4+ immunonegative SAN cells. We focused on to 3D-relationship between pacemaker meshwork and neuronal plexus.

**Fig. 3.**
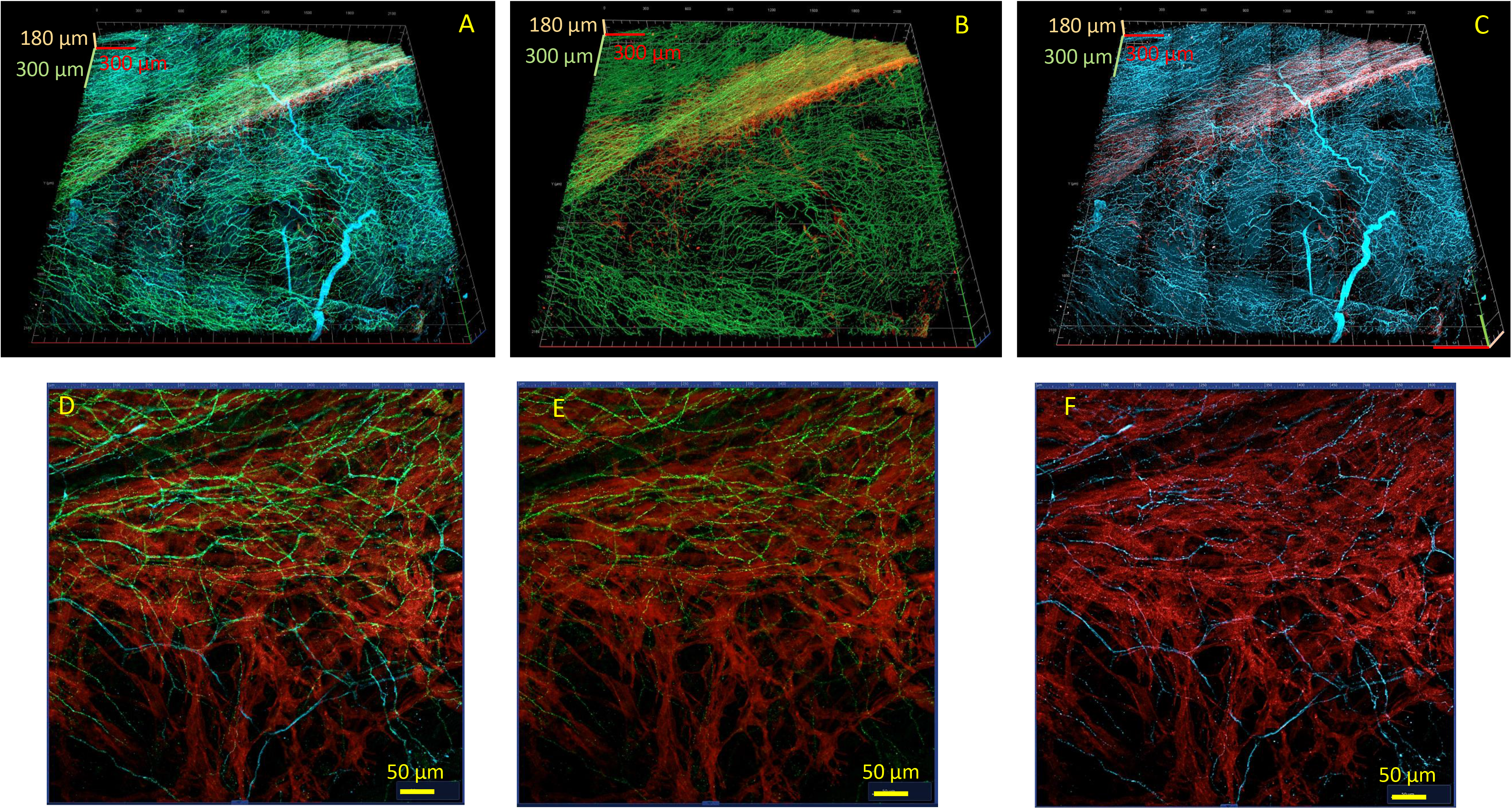
3D-image of the cholinergic and adrenergic neuronal plexus within the central part of the SAN and adjacent areas tiled from Z-stacks of 2D images (400µm by 400µm) that have been recorded with confocal microscopy by optical slicing of the whole-mount heart preparation. Panoramic 3D-images showed all immunoreactive fibers from endocardial to epicardial side through 300µm of depth. **Panels A, B and C** illustrate comprehensive 3D-images of the density and fine structure of the plexus, rich in both cholinergic and adrenergic sub-plexuses, dispersed around the HCN4^+^ immunoreactive area (**red color**). These panoramic 3D-images covered an area of 2 mm by 2mm. **Panel A** illustrates a 3D-image of the neuronal plexus of VAChT (**green**) and TH (**cyan**) immunoreactive neuronal fibers within SAN and in adjacent areas. The crista terminalis lies along the border of the HCN4+ immunoreactive meshwork (**red**) towards the top of the image and the interatrial septum is located at the bottom of the image. The 3D pattern of cholinergic innervation of SAN tissue (**Panel B**) did not resemble the 3D pattern of adrenergic innervation (**Panel C**). **Panel D** depicts both autonomic plexuses innervating the HCN4+ network. **Panels E, F, G** represent the same 2D- optical slice illustrating HCN4^+^ immunoreactive pacemaker cells (**red**) with VAChT+ (**green**) cholinergic and TH+ (**cyan**) adrenergic neuronal fibers that showed different patterns of HCN4^+^ meshwork innervation within the same slice. Although cholinergic neurites (**Panel F**) did not innervate the lower part HCN4+ meshwork, they exhibited higher density in the upper part than adrenergic neurites, which were detected across the entire image (**Panel G**).

Triple immunolabelling of the SAN with VAChT, TH, and HCN4 showed a comprehensive, dense plexus of fine nerve fibers around pacemaker cells in all sites of the HCN4-positive meshwork, spanning both the SAN and pectinate muscles of the right and left atria. Neuronal fibers of the SAN plexus also extended to cover contiguous areas along the borders of the HCN4+ meshwork. Neural plexus density was noticeably greater within the HCN4-positive pacemaker meshwork and adjacent areas than in the surrounding atrial walls, in accordance with previous reports (18, 19).

SAN 3D images at millimeter scale in **Fig 3. A-C** showed that 3D-patterns of cholinergic and adrenergic innervation were irregular and did not resemble each other. The 3D density patterns of adrenergic and cholinergic fibers was markedly patchy and created areas with dominant cholinergic innervation. In **Fig 3B**, a group of cholinergic fibers creates a chord of dense neuronal fibers that passes through the “body” of the HCN4+ pacemaker cell meshwork (**red**) from the upper right side of the picture to the lower left. Notably, while the adrenergic plexus showed homogeneous innervation across the SAN (**Fig. 3C**), there were portions of the pacemaker cell meshwork with an apparent scarcity of cholinergic neurites (**Fig. 3 and Fig 4**). **Panels D, E, and F** show the innervation within the central SAN pacemaker cell meshwork at higher resolution in 2D images taken from the 3D z-stacks.

**Fig. 4.**
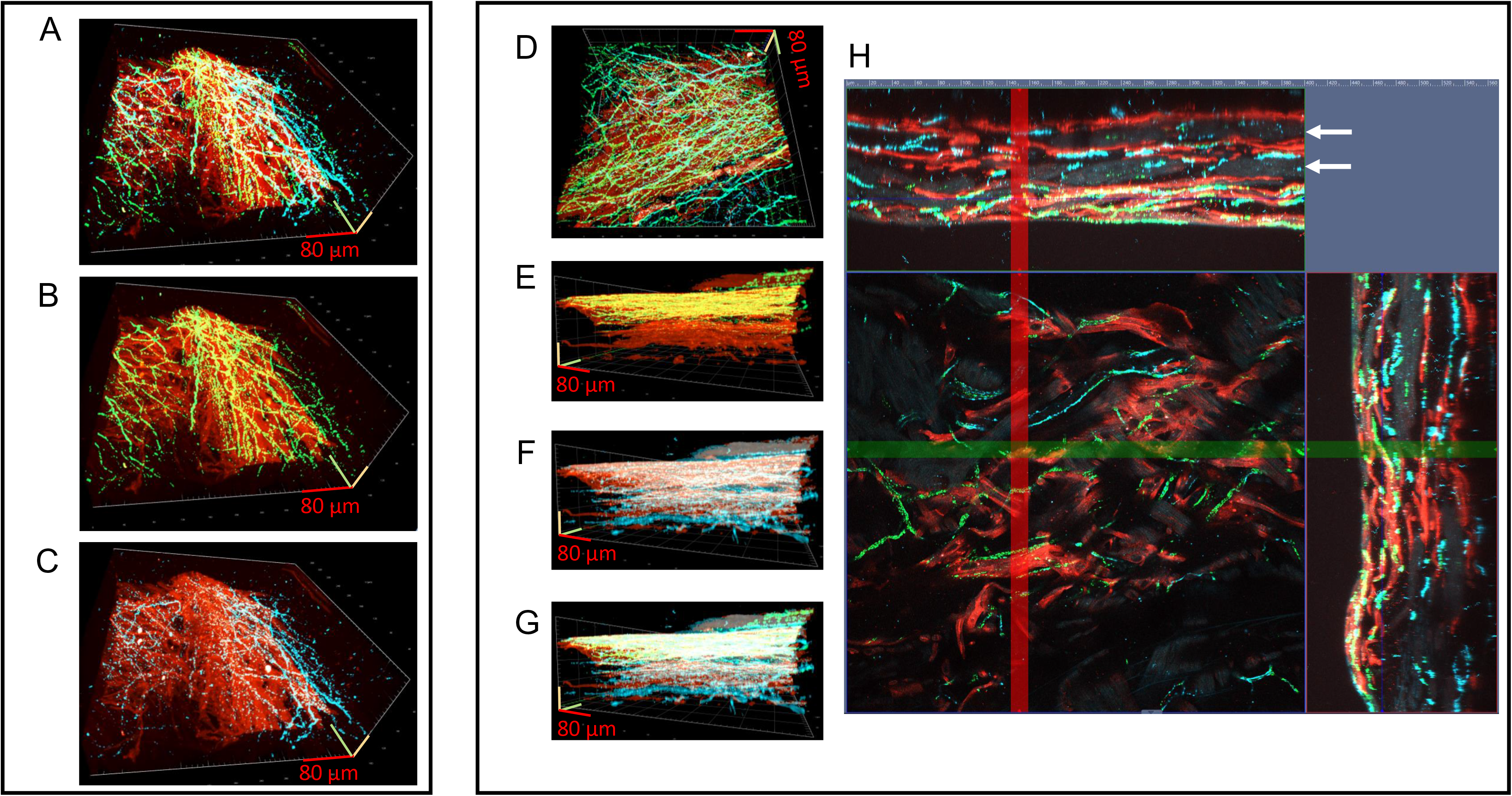
3D-images of SAN tissue of 400μm by 400μm and by 150μm illustrate 3D-density gradients of the cholinergic and adrenergic neuronal plexus enwrapping HCN4+ meshwork. This region of SAN tissue had 3 layers of HCN4+ immunoreactive cells (**red color**): one thick multicellular layer on the endocardial side and two monolayers, one in the middle and one on the epicardial side. Two “sleeves” of HCN4-immunonegative SAN cells, detected by background fluorescence, between layers of HCN4-immunopositive pacemaker cells created the 3D-cytoarchitecture of the stratified tissue. **Panel A** shows a 3D-image of SAN tissue, seen from the endocardial side, with triple immunolabelling of the VAChT (**green**) and TH (**cyan**) immunoreactive neuronal plexus unevenly distributed across the XY axes of the HCN4+ immunoreactive (**red**) meshwork. This 3D-image of the TH and VAChT immunoreactive neuronal plexuses highlights the dominance of cholinergic innervation in the central region of the HCN+ immunoreactive meshwork shown. **Panel B** illustrates the cholinergic neuronal plexus, which exhibited an even 3D-innervation pattern. **Panel C** illustrates the adrenergic neuronal plexus, with barely detectable neuronal fibers in the centre of the image. **Panel D** illustrates the 3D-image of the neuronal plexus seen from endocardial to epicardial side reconstructed from a series of Z-stack images taken within central part of SAN. The plexus of combined cholinergic (**VAChT-green**) and adrenergic (**TH-cyan**) fibers did not show a 3D-gradient across the XY plane of the HCN4-immunoreactive (**red**) pacemaker cell meshwork. **Panel E** shows a 3D graphical analysis of the Z-stack, with the endocardial (top)-endocardial (bottom) gradient of cholinergic innervation of the HCN4+ meshwork. Out of the 150μm depth of the HCN4+ meshwork, 70μm, from the endocardium to the mid SAN, were innervated by cholinergic neurites. **Panel F** shows a graphical analysis of the adrenergic neuronal fibers that innervated the HCN4+ meshwork evenly from the endocardial (top) to the epicardial (bottom) side. **Panel G** illustrates superimposed adrenergic and cholinergic innervation, which together create a gradient of predominantly excitatory innervation within the HCN4+ meshwork. **Panel H** illustrates virtual slicing of the Z-stack from epi-to-endocardial side at X and Y axes, shown by red (Y axis) and green (X axis) bands. The central image shows a 2D-picture within the Z-stack. The upper and right images are reconstructed virtual slices, approximately 150μm thick from epicardial to endocardial side, at the X and Y axes. Virtual cuts highlighted cholinergic and adrenergic varicosities (green and cyan dots correspondingly) within neurites in close proximity to HCN4+ membranes and confirmed that cholinergic fibers innervated pacemaker cells in half of the tissue from the endocardial side. “Sleeves” of HCN4 immunonegative tissue are indicated by **white arrows**.

Cholinergic neurites were mostly absent in the lower part of the HCN4-meshwork, as illustrated in **Figure 3 Panel E**, but showed overall higher density in the innervated 300um by 300um area than adrenergic neurites, which were present across the whole 2D image (**Fig. 3. Panels D, E, F).** Thus, the pacemaker meshwork is densely innervated, with uneven distribution of cholinergic and adrenergic projections.

**Figure 4** shows 3D-images of an area of SAN tissue 400μm by 400μm and by 160μm in depth, that exposes the fine details of density gradients of the neuronal plexus enwrapping HCN4+ meshwork (**Fig. 4**). 3D-imaging of the SAN tissue with triple immunolabelling showed that VAChT and TH immunoreactive fibers were unevenly abundant across XY axes of the HCN4+ immunoreactive meshwork. Superimposed 3D-images of the TH and VAChT immunoreactive neuronal plexus showed area of the HCN+ immunoreactive cellular meshwork in the center of the 3D-image, which was recorded from the epicardial side (**Fig. 4A**) with the dominant cholinergic innervation. As previously mentioned, cholinergic fibers had homogeneous density (**Fig. 4B**). Adrenergic innervation had high density in the right side of the 3D image, towards the crista terminalis; exhibited lower density on the left side, towards the interatrial septum; and was barely detectable in the central part of this high-zoom 3D image (**Fig. 4C**).

3D-imaging of the autonomic plexus in the “body” of the SAN, reconstructed from endocardial to the epicardial side, did not show any 3D-gradient across the XY plane though the HCN4-immunoreactive pacemaker cells meshwork (**Fig. 4D**). However, 3D visualization of the SAN tissue found endo-epicardial gradient of HCN4+ immunoreactive meshwork cholinergic innervation in which half of the tissue, from the middle to the epicardium, lacked cholinergic fibers (**Fig. 4E**). Cholinergic neurites innervated the HCN4+ meshwork up to a depth of 70μm from the endocardial side, out of the 150μm total Z-depth of the right atrium. Adrenergic neuronal fibers innervated the HCN4+ meshwork evenly from the endocardial to epicardial side (**Fig. 4F**). Combined adrenergic and cholinergic innervation patterns created an area of dominant excitatory input within the HCN4+ meshwork. We may theorize this to be a functionally significant region in which pacing frequency may be increased without cholinergic inhibition, while in adjacent parts of the meshwork adrenergic action will be counteracted by acetylcholine release from cholinergic synapses.

The side and the top views of **Figure 4, Panels A to G** show the overlap of HCN4+ immunoreactive cells and neuronal fibers on top of each other, in a Z-stack that shows all immunoreactive cells detected within it. This overlapping of cells and fibers masked the fine details of the tissue 3D-cytoarchitecture. This issue was addressed by virtual slicing of the Z-stack from epi-to-endocardial side at any position of X or Y axes (**Fig. 4 Panel H** position of X and Y axes are shown by green and red band correspondingly). The central image of the panel H in the Figure 4 show an example of a 2D-image from the series of scans taken to reconstruct the Z-stack. The upper and right images represent virtual slices of the approximately 150μm thick Z-stack, seen from epicardial to endocardial side, at the position indicated by the X and Y axes.

The green and cyan dots seen in **Figure 4, Panel H** are cholinergic and adrenergic varicosities within neurites near HCN4+ membranes. The distribution of varicosities from endocardial to epicardial side confirmed that cholinergic fibers innervated the SAN within an area extending approximately half the distance from endocardial to the epicardial side.

This region of SAN tissue had 3 layers of HCN4+ immunoreactive cells: one thick multicellular layer on the endocardial side, and two monolayers, one in the midwall and one beneath the epicardium, that were separated by two strata of HCN4-immunonegative SAN cells, inferred by background fluorescence. Scanning of the Z-stack with virtual cuts showed the SAN to be defined by interweaving layers of HCN4 positive and negative cells (**Figs. 4, 5**).

**Fig. 5.**
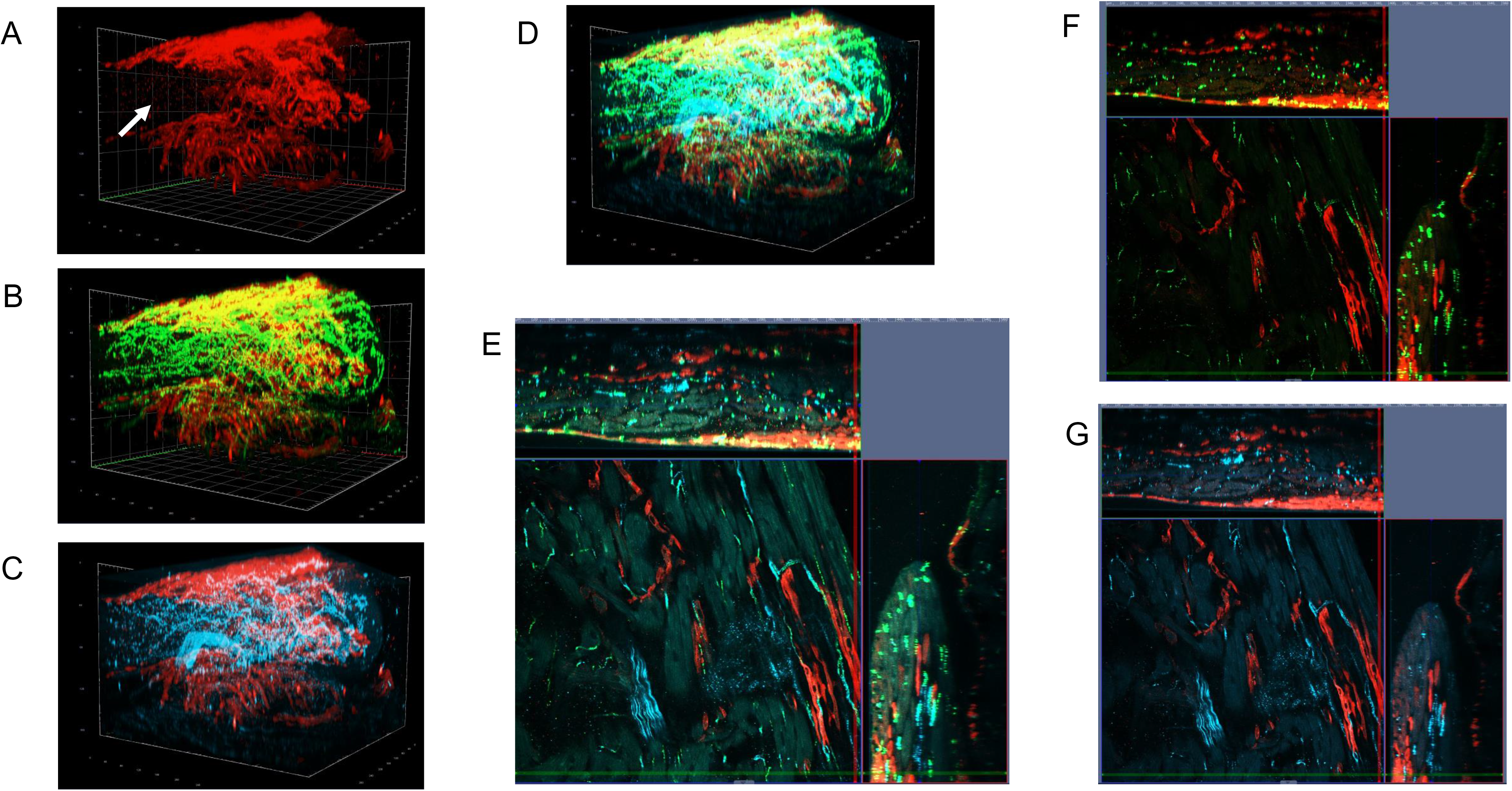
3D-images of SAN tissue, 400μm by 400μm and by 150μm, illustrating the 3D-cyroarchitecture of the HCN4-immunoreactive meshwork and density gradients of the cholinergic and adrenergic neuronal plexuses. **Panel A** illustrates the 3D-cytoarchitechture of the HCN4-immunoreactive (**red color**) cells in the SAN region adjacent to the crista terminalis. Cells from the crista terminalis folded to make a “cave” (**white arrow**) out of the HCN4+ meshwork, creating a 3D “dent-like” cytoarchitecture. The “sleeves” of HCN+ cells partially enveloped the top and the bottom of the crista terminalis. The empty invagination in the middle of the 3D-image of the HCN4+ meshwork is where cells from the crista terminalis are folded into the meshwork. Two layers HCN4-immunoreactive cells on endocardial and epicardial side were connected by an intermediate layer. HCN4+ immunoreactive cells had maximum density of **##** on the endocardial side (top), which gradually decreased towards the epicardial side (bottom) of the 3D-image. **Panel B:** 3D-image showing the adrenergic neuronal plexus (**TH-cyan color**) evenly innervating the central part of the HCN4+ meshwork (**HCN4-red color**) and cells within the crista terminalis (empty space within the HCN4 “cave”). **Panel C** 3D-image illustrates the gradient of cholinergic neural plexus (**VAChT-green**), which was highly dense within the HCN4+ meshwork (**HCN4-red color**) on the endocardial side but had decreased presence towards epicardial side. **Panel D** The side view of the 3D-iamge illustrates the endo-to-epicardial gradient of both cholinergic (**VAChT-green**) and adrenergic (**TH-cyan**) neuronal plexus. Both adrenergic and cholinergic fibers richly innervate the central part of the HCN4-immunoreactive meshwork and crista terminalis. **Panels E, F, G:** Each panel has a central 2D-picture as well as upper and left images, illustrating triple immunolabelling of the SAN with **HCN4 (red), VAChT (green)** and **TH (cyan)** antibodies. The central 2D-picture taken from the Z-stack series illustrates single HCN4+ pacemaker cells penetrating between HCN4-immunonegative cells within the “cave” shown in panel A. The upper and right 2D-images illustrate reconstructed virtual cuts (approximately 150μm in thickness) taken from the endocardial part (top) to the epicardial part (bottom), through X (**upper 2D-image**) and Y axes (**right 2D-image**). “Sleeves” from the HCN4+ meshwork at the endocardial side (bottom of the image) thinned in the direction of atrium. The multilayer of HCN4+ cells partially covered the crista terminalis from the epicardial side. **Panel F** illustrates cholinergic varicosities within neurites of the plexus which richly innervated HCN4-immunoreactive cells in the endocardial side and HCN4-immunonegative cells within the “cave”. **Panel G** illustrates adrenergic varicosities that were abundant among cells within the “cave” and were barely detectable near HCN4+ pacemaker cells on the endocardial and epicardial sides.

3D-imaging of portions of the HCN4-meshwork adjacent to the crista terminalis showed the intertwining of HCN4+ immunoreactive pacemaker and immunonegative SAN cells. The crista terminalis (**white arrow**) was partially enveloped at the top and bottom by the HCN4+ meshwork (**Fig. 5A**).

The side panels in **Figure 5** illustrate the endo-epicardial distribution of cholinergic and adrenergic neuronal plexus (**Fig. 5D**). Interestingly, adrenergic neuronal fibers innervated evenly the central part of the HCN4+ meshwork and cells within the crista terminalis (**Fig. 5C**), while cholinergic nerves had higher density on the endocardial side that decreased towards epicardial side (**Fig. 5B**). Remarkably, both cholinergic and adrenergic innervation had similar density within the HCN4-immunonegative “cave” and within the central region of the HCN4+meshwork.

The cross-sectional 2D-images from the Z-stack in **Figure 5** (thickness of approximately **150μm**) showed that HCN4+ pacemaker cells penetrated between HCN4-immunonegative SAN cells from the Crista Terminalis (**Fig. 5E**). The **upper 2D-panels in Fig. 5E, F, G** demonstrate that the HCN4+ meshwork “sleeve” at the endocardial side (bottom of the panels) thinned in the direction of the atrium. Cholinergic varicosities within neurites of the plexus richly innervated HCN4-immunoreactive cells in the endocardial side and HCN4-immunonegative cells within the “cave” (**green dots in Fig. 5F**). Adrenergic varicosities were abundant among cells within the “cave”, and were barely detectable near HCN4+ pacemaker cells on the endocardial and epicardial sides (**cyan dots in Fig. 5G**).

### Triple immunostaining of Pacemaker and Peripheral Glial Cells in the SAN with GFAP, S100B and HCN4 antibodies

3D-images of the SAN with triple immunolabelling of adrenergic, cholinergic nerves and HCN4-pacemaker cells showed that the neural fibers density is comparable to that of pacemaker cells within the meshwork. However, these images did not include one major player: peripheral glial cells (**PGCs**). Peripheral glial cells have been characterized within the mesenteric plexus by double immunostaining (21), but have never been investigated within the SAN at the macroscopic level. Since the early 1970s glial fibrillary acidic protein (GFAP), an intermediate filament protein, has been accepted as a predominant component of the 10-nm intermediate filaments of astrocytes (22). Although GFAP may be expressed by several cell types, co-expression of Ca^2+^ binding protein S100B defines a restricted population of cells, which includes non-myelin-forming Schwann cells (**NMFSc**), enteric glia (**EGc**) and astrocytes (21, 23). GFAP and S100B, in combination with morphological characteristics, have both been used for the identification of peripheral glial cells (**PGCs**).

We designed a triple immunostaining procedure (S100B, GFAP, and HCN4) to identify peripheral glial cells (**PGCs**) and SAN pacemaker cells simultaneously. The panoramic 2D-image (4mm by 1.2mm) in **Figure 6A,** tiled from series of images, shows the HCN4-immunoreactive meshwork in which the head, the body, and tail could be clearly distinguished. The “head” of the meshwork located in the right side of the image, near the superior vena cava, continued to the “body” in the center and “tail” of the meshwork on the left side of image near the inferior vena cava. We found cells immunoreactive to GFAP and S100B with a somatic shape resembling “stars” scattered among HCN4-immunoreactive cells, across the area of the SAN GFAP and S100B immunoreactive somata of peripheral glial cells (PGCs) embedded within the SAN were interconnected by GFAP+ branches and created a “web”-like regular pattern within and outside the HCN4^+^ immunoreactive meshwork. This web of PGCs within the SAN has not been imaged previously.

**Fig. 6.**
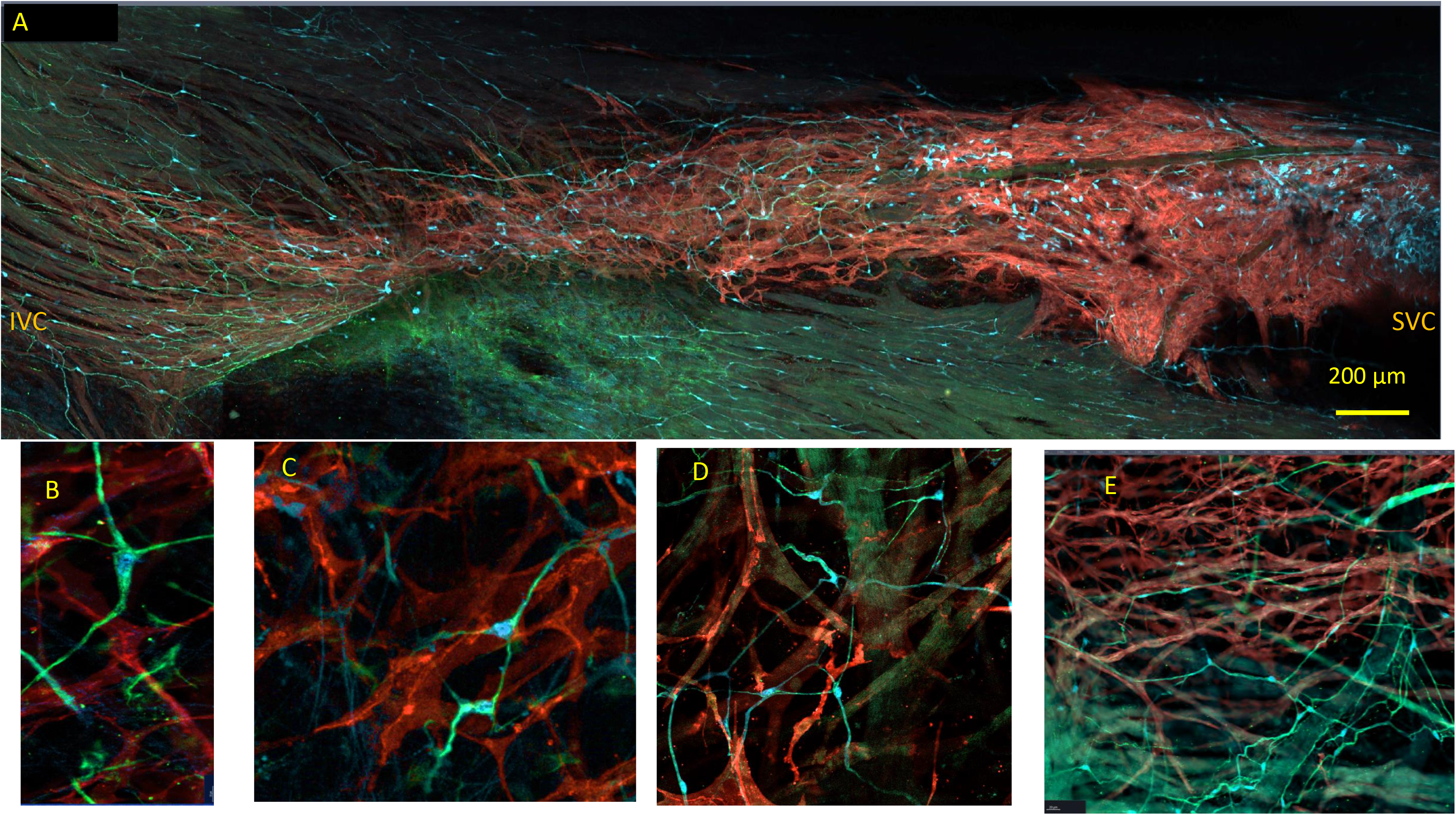
2D-images of the whole-mount SAN preparations with triple immunolabelling illustrate peripheral glial cells (**PGCs**) and SAN pacemaker cells imaged by optical slicing. All “star” shaped PGCs detected within the SAN expressed both GFAP (**green color**) and S100B (**cyan color**). **Panel A:** Panoramic 2D-image (4mm by 1.2mm), tiled from a series of images showing the 2D cytoarchitecture of the HCN4-immunoreactive meshwork with the “head” of the meshwork located in the right side of the image, near the superior vena cava, and the “body” in the center which thinned and enlarged to the “tail” of the meshwork on the left side of image near the inferior vena cava. Peripheral glial cells are scattered between HCN4-immunoreactive cells (**red color**), across the SAN. GFAP+ and S100B+ PGGs embedded within the SAN were connected by GFAP+ branches and created a web within and outside the HCN4-immunoreactive meshwork that exhibited a regular pattern. Other cellular clusters located in the area of the “head” and “body” of the HCN4-immunoreactive meshwork were not connected with GFAP+ branches. **Panels B, C, D and E** illustrate peripheral glial cells (PGCs) immunoreactive to GFAP (**green**) and to S100B (**cyan**) among HCN4-immunoreactive cells (**red**) imaged with high optical magnification. GFAP+ was detected in higher levels than S100B+ within the branches PGCs. **Panel E** illustrates the web of PGCs that made contact with the walls of the blood vessels within SAN. Somata and processes of PGCs were closely attached to the endothelial cells outlining the lumen of the blood vessels within the tissue.

A closer look to the PGCs within the SAN showed “star” shaped cells with elongated soma of 10.2+0.8μm, which had 2, 3, 4 or 5 projections (**Fig. 6 B, C, D**). Those PGCs found to express **both** S100B and GFAP exhibited uniform morphology. Interestingly, the web of PGCs was found to also surround the borders of the lumen of blood vessels (**Fig. 6 E**).

#### S100B+/GFAP- interstitial cells within the SAN

We found clusters and individual examples of S100B+ cells that were GFAP immunonegative, with the cellular morphology sharply different from the “star” like cellular shape of PGCs. 3D-images of the triple immunolabelled whole mount SAN preparation showed that this new type of cell was found within the borders of the SAN (**Fig. 7**). The 3D-image of S100B-immunoreative and GFAP immunonegative cells, built up by tiled series of Z-stack images, confirmed that S100B+/GFAP- cells were detected to be mostly localized within the HCN4-immunoreactive pacemaker meshwork (n=5). Scattered somata of S100B+/GFAP- cells exhibited an irregular pattern of cell clustering and distribution, unique for each immunolabelled SAN preparation. For example, S100B+/GFAP- cells shown in the 3D-panoramic image (**Fig. 7**) are grouped in one cluster in the “head” of HCN4+ meshwork (right side of the 3D-image near SVC) and a second cluster on the left, towards the IVC. Some of the S100B+/GFAP- cells were scattered across the “body” of the HCN4+ meshwork, and a couple of small clusters can be detected in the “tail” section of the SAN (left side of the image near the IVC). In another SAN preparation, 60-80% of detected cells formed two clusters near the SVC and the IVC. S100B+/GFAP- cells were scattered, without a noticeable pattern, across the whole HCN4^+^ meshwork in another two SAN preparations. Four of the whole mount SAN preparations had a mix of clustered and scattered S100B+/GFAP-cells. The number of S100B+/GFAP- cells within SANs, as detected by counting DAPI-labeled nuclei, varied from 21 to 148, giving in average 72.3+18.6 of cells (n=7). It should be noted that it is difficult to exactly quantify S100B+/GFAP- cells because they often interlace and thus obfuscate their total number, further work using more detailed techniques would be needed in future studies to determine their number in relation to other cell types within the SAN.

**Fig. 7.**
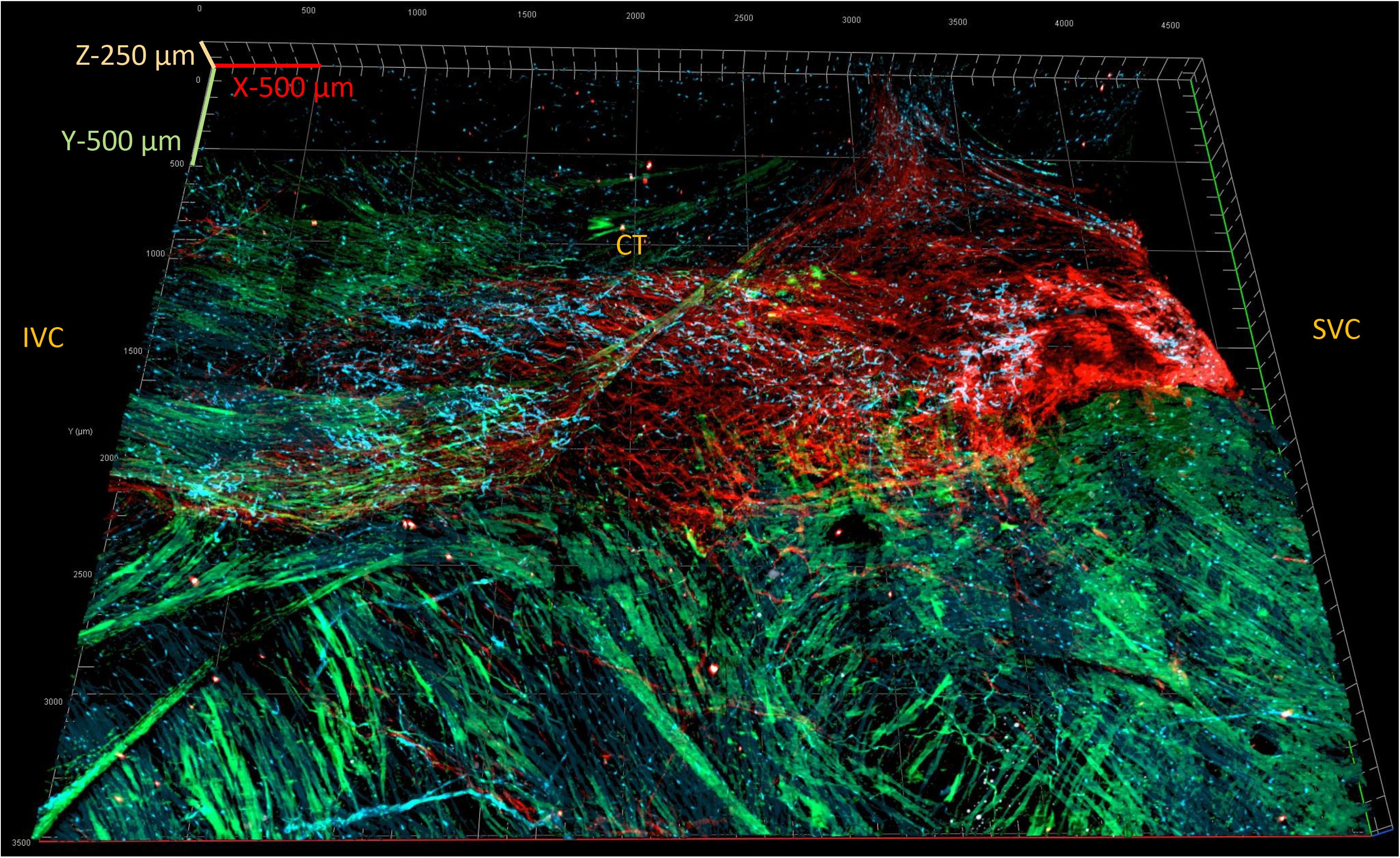
3D-image of the whole-mount SAN preparation shows an area 4.5mm long, 3.5mm wide, and 250μm deep. Triple immunolabeling and 3D reconstruction showed a new type of S100B^+^ (cyan)/GFAP^-^ (green) cells detected within and near the HCN4-immunoreactive meshwork (red color). The “head” of the HCN4-immunoreactive meshwork is on the right side of the image, near the superior vena cava (SVC). Part of the right auricle is shown in the image, above the crista terminalis (CT). The tail of the HCN4-immunoreactive meshwork is extended to the inferior vena cava (IVC). The auricle lacks S100B^+^ (cyan)/GFAP^-^ (green) cells.

S100B+/GFAP- cells were situated between, and in close proximity to, HCN4-immunoreactive pacemaker cells. By definition, this classifies them as interstitial cells of the sinoatrial node. The geometry of their somata and cellular extensions, as well as the lack of expression of GFAP protein, undoubtedly separated S100B+/GFAP- interstitial cells from the web of S100B+/GFAP+ peripheral glial cells (**Fig. 8**). The pattern of S100B+/GFAP- processes of interstitial cells was distinct from the web of GFAP+/S100B+ glial cell extensions interconnecting PGCs somata, even within areas of the SAN in which both cell types intertwined (**Fig. 8A**). Further, while the pattern of PGC branches across the HCN4+ immunoreactive meshwork was regular (i.e., forming a web), that of S100B+/GFAP- intercellular connections did not conform to any distinguishable order (**Fig. 8 A, B, C, D**). PGCs radiated GFAP+ branches that shadowed neuronal fibers, while we did not detect S100B+/GFAP- interstitial cells enwrapping neurites from SAN neuronal plexus. Two images shown in **Figure 8** illustrate radiating branches of adrenergic (**Fig. 8E**) and cholinergic fibers (**Fig. 8F**) that resembled the web of GFAP+ processes (**Fig. 8 A, B, C, D**). The S100B+ somata of PGCs were often found within bundles of adrenergic (**Fig. 8E**) or cholinergic (**Fig. 8F**) varicosities. S100B^+^/GFAP- interstitial cells were exclusively found outside of nerve bundles or varicosities.

**Fig. 8.**
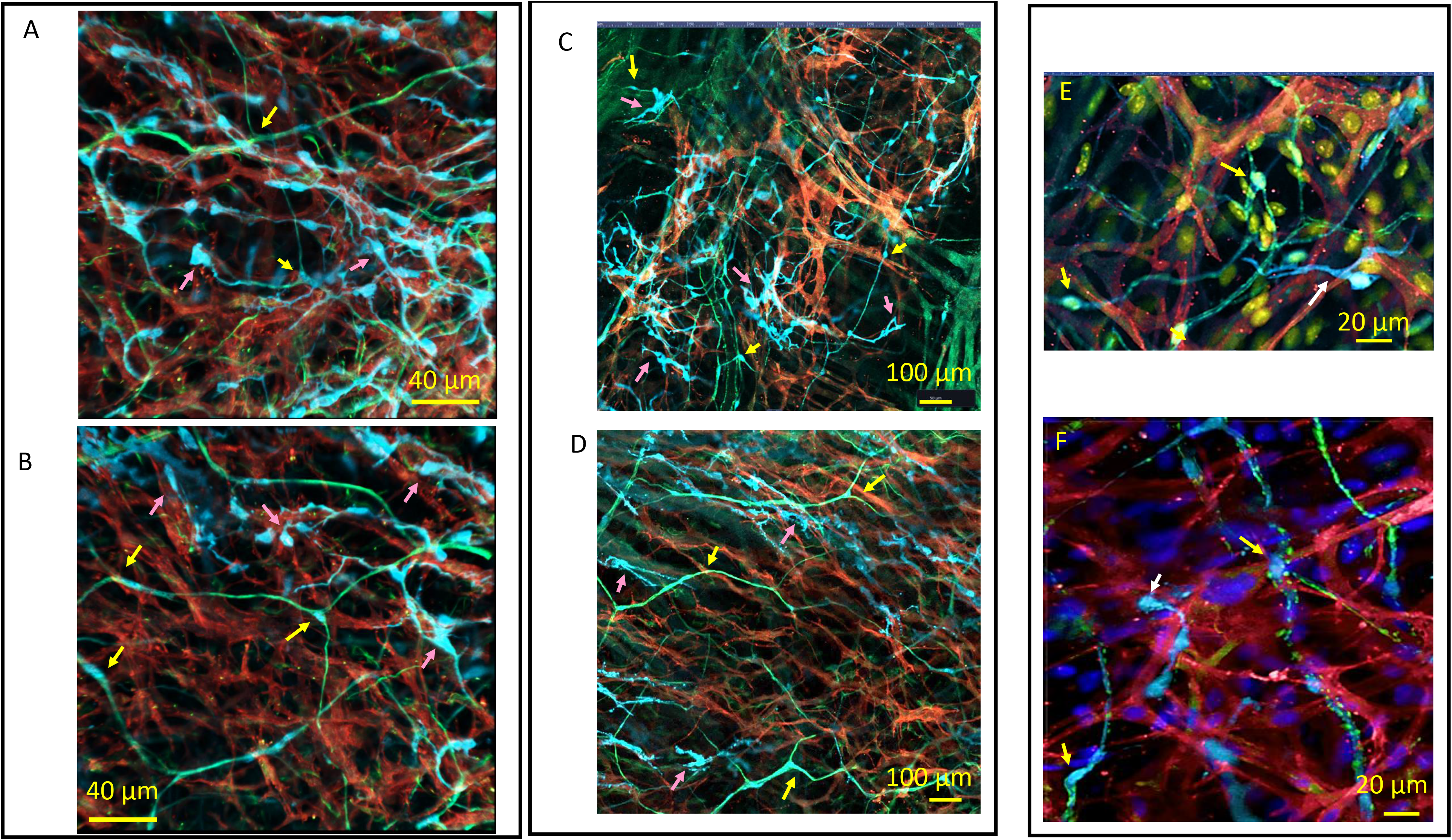
2D-images of S100B+/GFAP- interstitial cells and peripheral glial cells embedded within the HCN4-immunoreactive meshwork, imaged from whole-mount SAN preparations. Examples of S100B+/GFAP-interstitial cells are indicated by pink arrows. The S100B+ somata of PGCs, indicated by yellow arrows, were detected within adrenergic and cholinergic fibers. TH and VAChT immunoreactive varicosities surrounded the S100B+ somata of PGCs, while S100B+ interstitial cells were only found outside them. **Panels A, B, C and D** illustrate S100B+/GFAP-interstitial cells (**cyan color**) lying between, and in close proximity to, HCN4-immunoreactive pacemaker cells (**red color**). The regular web of long extensions from GFAP and S100B immunoreactive peripheral glial cells (**green color**), indicated by yellow arrows, does not overlap with the GFAP-immunonegative processes of S100B+/GFAP-interstitial cells, even within intertwined clusters of cells. Higher levels of GFAP than S100B were detected in the extensions of peripheral glial cells. The recognizable web of glial cells exhibits a pattern different from the irregularly scattered S100B+/GFAP-branches. **Panel E** shows a 2D-image of the radiating branches of adrenergic TH+ (**green**) fibers as well as S100B+ interstitial cells (**cyan**), among HCN4-immunoreactive pacemaker cells (**red**). **Panel F** shows a 2D-image of the radiating branches of cholinergic VAChT+ (**green**) fibers as well as S100B+ interstitial cells (**cyan**), among HCN4-immunoreactive pacemaker cells (**red**). The pattern of neuronal fibers resembles the web of GFAP+ processes from PGCs.

### Anatomical interaction between HCN4+ pacemaker cells and S100B+/GFAP- interstitial cells

The S100B^+^/GFAP^-^-SAN interstitial cells and HCN4^+^ pacemaker cells interact anatomically in groups and clusters. We characterized instances of anatomical interaction between S100B+cells and HCN4+ cells by the shape of cellular branches and geometry of the endings of cellular extensions. S100B+/GFAP- cells showed both tapering and dilating extensions, which changed diameter right after exiting the soma (**Fig. 9**). These extensions could broaden and narrow along their length. We defined these long tapering cellular extensions that flattened along the surface of a target cell as “spiculum”-type. These were not observed within the web of S100B+/GFAP+ PGCs, in which cellular extensions had a constant diameter and did not show sinuses or cytoplasmic dilations along their branches. The S100B+ interstitial cells shown in **Figure 9**, **panel A**, extended spicula of 78μm alongside HCN4+ pacemaker cell so close that extracellular space between two cells could not be detected with the optical resolution of the confocal microscopy we employed. Spicula from the cluster of S100B+ cells shown in **Figure 9A** targeted and adhered to the same group of HCN4+ cells. Spicula could bifurcate into multiple distinct tapering processes and wind around a target HCN4+ immunoreactive cell. Anatomical connections between S100B+ interstitial cells and HCN4+ cells by spicula were defined as fibrous “cotton” type interactions.

**Fig. 9.**
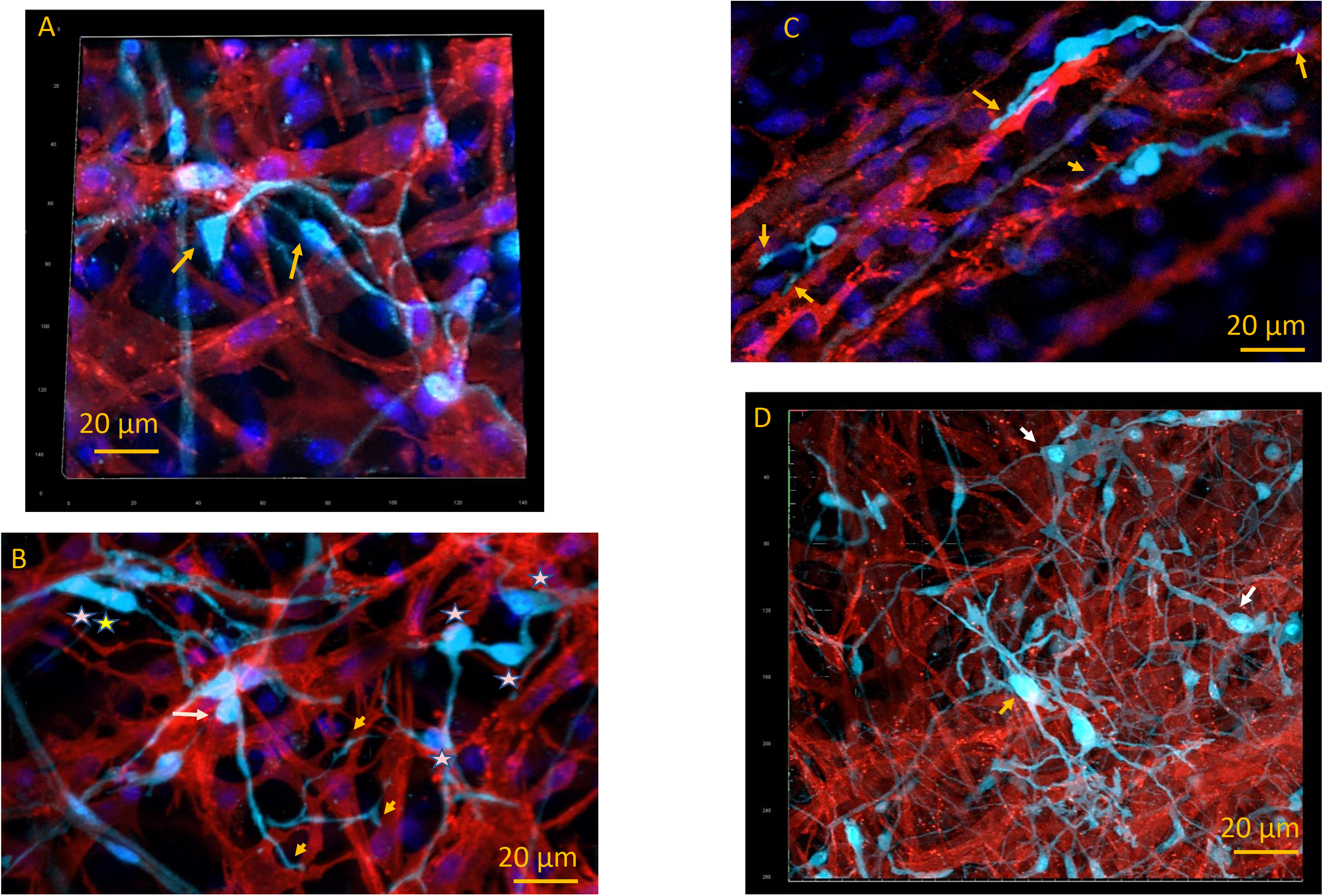
Fibrous “cotton” type of anatomical interaction between S100B^+^ immunoreactive interstitial cells and HCN4^+^ pacemaker cells. **Panel A:** 3D image (15 μm thick) illustrates two unipolar S100B+ cells (**cyan color**) projecting tapered spicula that bifurcated on the HCN4-immunoreactive cells (**red color**). These spicula adhered so close to the HCN4+ cell that extracellular space could not be detected by optical confocal microscopy. DAPI staining highlights nuclei (**blue color**). The soma of the unipolar S100B+ cell, and the bifurcations of their spicula are indicated by yellow arrows. **Panel B:** 2D-image that illustrates unipolar S100B+ cell, indicated by yellow arrow, connected to several HCN4+ pacemaker cells by one bifurcating spiculum. Groups of cells (**yellow star**) or clusters of S100B+ somata (**2 yellow stars**) attached to HCN4^+^ cells and were interconnected by short extensions in a “nodal”-like net cytoarchitecture. S100B+ cells from this “nodal”-like net extended long spicula to adjacent HCN4+ cells. **Panel C:** 2D image that illustrates the spiculum of an S100B+ cell (**cyan**) that dilated in an “endfoot”-like structure, indicated by an arrow, adhering to HCN4+ pacemaker cells (**red**). The two pacemaker cells interconnected by one bipolar S100B+ interstitial cell also have a point of direct contact. **Panel D:** 3D image, 20μm thick, that illustrates composite fibrous “cotton” connections that include the spicula, “endfeet”, and “nodal”-like net of S100B+ (**cyan color**) cells within the meshwork of HCN4+ cells (**red color**).

Some HCN4+ pacemaker cells were connected to multiple S100B^+^ interstitial cells at once, while the majority of HCN4+ cells lacked connections with S100B+ cells altogether. Alternatively, one S100B+ cell could project spicula to several HCN4+ pacemaker cells. Examples of such S100B+ cells are shown in **Figure 9, Panel B** and **Figure 11**, **Panel E.**

**Fig. 10.**
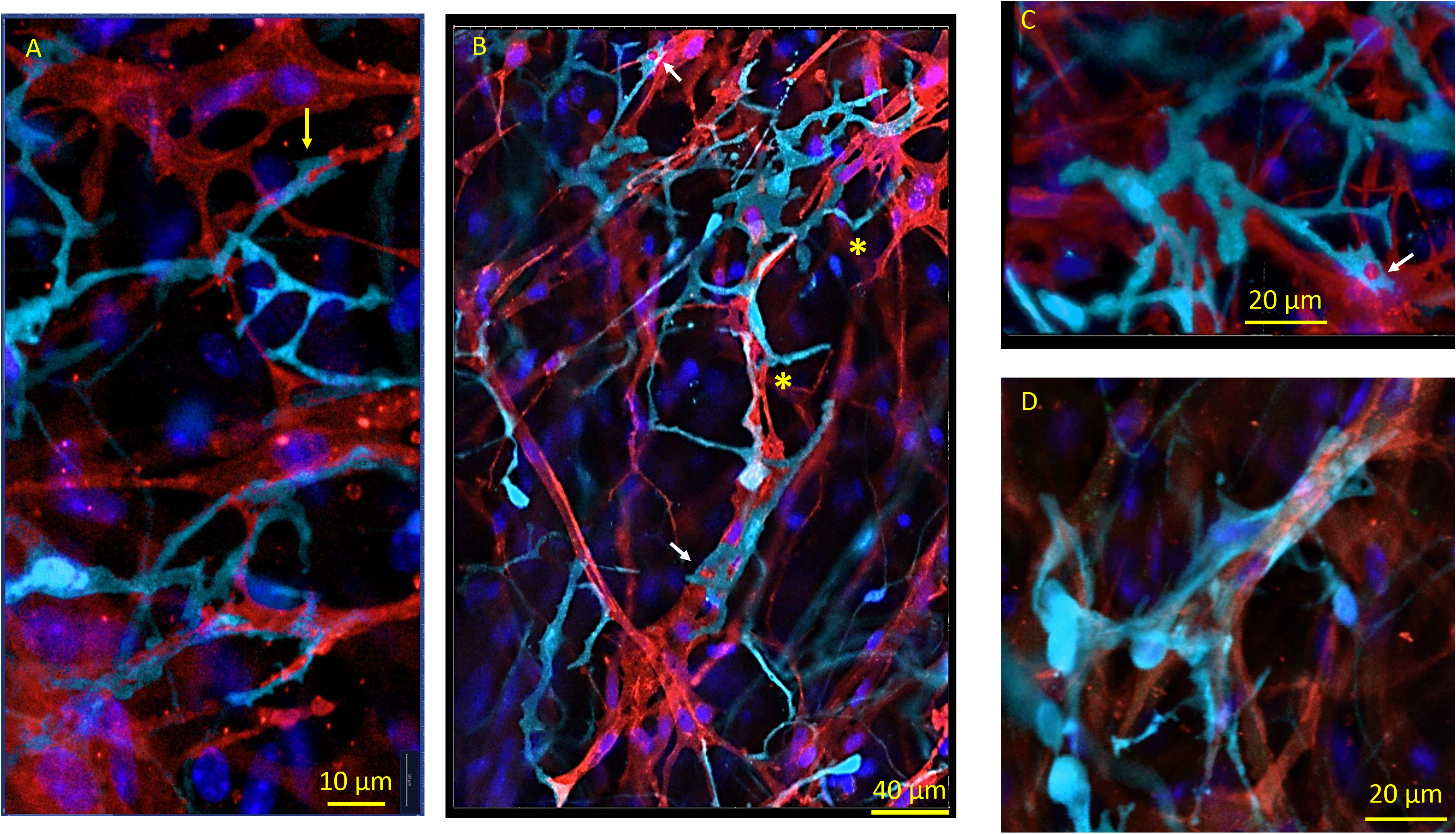
Anatomical interaction between amoeboid-like S100B-imunoreactive interstitial cells and HCN4+ pacemaker cells. **Panels A, B and C** illustrate amoeboid S100B+ interstitial cells (**cyan**) with flattened cellular extensions. Flattened S100B+ extensions, or pseudopodia, were 1-2 µm wide and manifested dilations. S100B+ pseudopodia could fold, changing the initial direction of their projection, or bifurcate and produce branches as indicated by yellow stars. The “plier”-like terminal dilation of an S100B^+^ pseudopodium, enclosing an appendage from an HCN4+ pacemaker cell (**red**), is indicated on Panel A by yellow arrow. The “patch”-like dilation of an S100B+ pseudopodium that encircled a “patch” of the membrane of an HCN4+ cell is indicated by a white arrow. **Panel D** shows S100B+immunoreactive cells enwrapping a group of HCN4+ pacemaker cells with a wide “ribbon”-like pseudopodium.

**Fig. 11.**
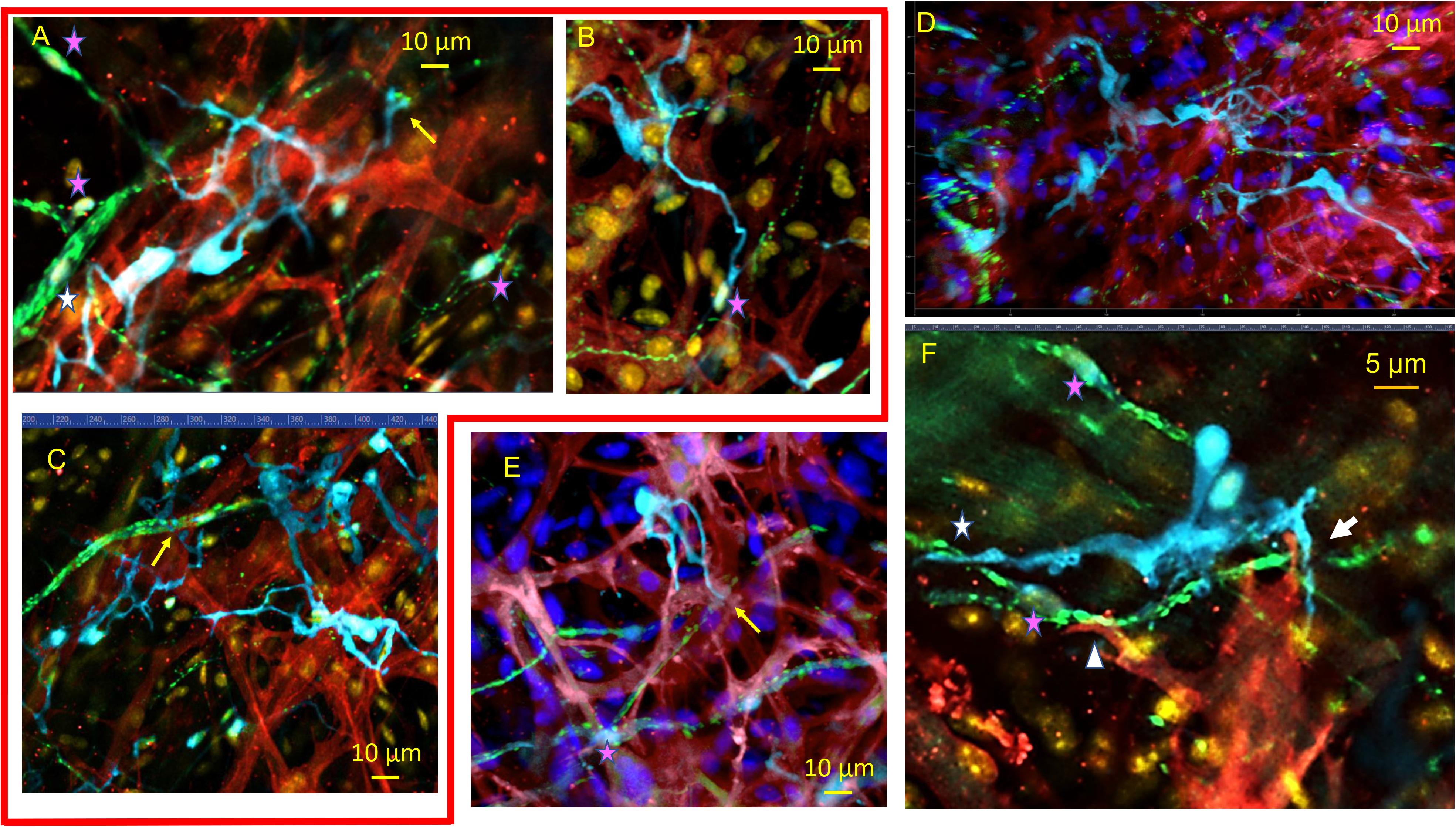
2D-images of a whole-mount preparation of SAN tissue with triple immunolabelling for S100B (**cyan**), HCN4 (**red**) and TH (**green**)/or VAChT (**green**). **Panels A, B, C:** 2D-images of SAN tissue with triple immunolabelling for S100B+ cells (**cyan**), HCN4-immunoreactive pacemaker cells (**red**) and TH+ adrenergic fibers (**green**) illustrate anatomical interactions between pacemaker cells, adrenergic nerves, and interstitial cells. **Panels D, E, F:** 2D-images of SAN tissue with triple immunolabelling for S100B+ cells (**cyan**), HCN4-immunoreactive pacemaker cells (**red**) and VAChT+ cholinergic fibers (**green**) illustrate anatomical relations between pacemaker cells, adrenergic nerves and interstitial cells. Pink stars on all panels indicate the nuclei of peripheral glial cells. S100B+ spicula extended from “octopus”-like cells in panels **A, C and E** ended on TH+ varicosities (A and C) or on VAChT+ varicosities (E). Yellow stars indicate the S100B+ “endfeet”. An adrenergic neurite on panel A, indicated by a white star, overlaps with the spiculum of an “octopus”-like S100B+ cell. An amoeboid-like S100B+ cell in the upper right conner of **panel C** receives adrenergic innervation. Images on **panels C** (adrenergic nerves) and **D** (cholinergic nerves) illustrate that within one cluster of S100B+ cells, some cells could receive nervous input while others might not. **Panel F** illustrates a composite point of contact between three cells: one S100B-immunoreacrtive cell, an HCN4-immunoreactive cell, and fibers from the neuronal plexus. A white arrow indicates the region where cellular extensions from an intertwined couple of S100B+ cells, an HCN4+ pacemaker cell, and a cholinergic neurite colocalize within 1 µm of each other, manifesting an anatomical unit between the three cell types. A cytoplasmic extension from HCN4-immunoreactive cell, resembling a dendritic spine, made contact with a folded amoeboid S100B+ pseudopodium. A white star indicates the point of contact between an S100B+ cell and a cholinergic nerve, while a white triangle marks the point of contact between a cholinergic nerve and an HCN4+ pacemaker cell.

S100B+ spicula might terminate into an “endfoot”-like structure that adhered to HCN4+ pacemaker cells (**Fig. 9C**). Note that the two pacemaker cells interconnected through one S100B+ interstitial cell in the image also had a point of direct contact to each other. The size of the S100B+ endfeet varied from 3.4+1.8μm to 14.9+1.3μm, and could differ between “spot” or “stretched” endfeet. Both types of endfeet were equally scattered within the HCN4+ meshwork.

Groups of S100B+ somata could be closely interconnected by short processes or wide cytoplasmic dilations (**stars in Fig. 9B, C, D**) rather than the thin projections described above, creating what we define as “nodal”-type cytoarchitecture. Cells within these S100B+ “nodal” groups could connect to adjacent pacemaker cells either through long extensions (**yellow arrows in Fig. 9B**) or by having their somata directly adjacent to HCN4+ cells (**white arrow in Fig. 9B**). Often, S100B+ interstitial cells created composite webs of “nodes”, interconnected by the aforementioned “spiculum” type extensions (**Fig 9D**).

Another possible anatomical interaction between S100B-imunoreactive interstitial cells and HCN4+ pacemaker cells was defined as “ameboid-flattened” type. These Interstitial cells resembled amoeba (**Fig. 10**). We defined flattened S100B+ extensions from this type of interstitial cell as a “pseudopodia”, also by morphological analogy with the morphology of the amoeba. Somata of ameboid S100B+ cells manifested round or elongated oval shapes. Typically, S100B+ pseudopodia were 1-3µm wide, and could manifest dilations with diameter reaching 5 µm, often as wide as the somata from which they originated (**Fig. 10A, B**). Pseudopodal extensions could change their initial direction of branching and create angles, as well as bifurcate (**Fig 10**).

S100B+ pseudopodal dilations could either adhere to the surface of HCN4+ pacemaker cells or enwrap an appendage extended from a pacemaker cell (**Fig. 10A**). Terminal cytoplasmic dilations of S100B+ immunoreactive pseudopodia manifested unique geometry. Some S100B^+^ pseudopodial terminal dilations had a shape similar to “pliers”, that enclosed an appendage from an HCN4+ pacemaker cell (**Fig. 10A**). A second type of contact point was “patch”-like dilation of S100B+ pseudopodium, which adhered on to the HCN4^+^ cell’s membrane (**Fig. 10B and C**). The distance between S100+ immunoreactive pseudopodia and HCN4+ pacemaker cells in the contact points were so close that we could not detect a space between them at the resolution of our confocal microscopy. Finally, S100B+ cells enwrapped one or several HCN4+ pacemaker cells with a wide “ribbon”-like pseudopodium. S100B+ cells with spiculae, as well as those with ameboid extensions both manifested uni- or multipolar branches.

### Anatomical interactions between HCN4+ pacemaker cells, S100B+immunoreactive interstitial cells and the neuronal plexus

In addition to anatomical contact between S100B+/GFAP- immunoreactive interstitial cells and HCN4+ pacemaker cells or peripheral glial cells, both cholinergic and adrenergic neurites have been found near S100B+ cells (**Fig. 11**). We did not find any specific pattern of innervation among S100B+ cells which could be related to the shape of the S100B+ cell, to the number and type of processes, or to the number of contacts between S100B+ cells and HCN4+ pacemaker cells.

Unipolar or multipolar S100B+ cell extended dendritic-like processes toward neurites, which ended on or near the neuronal fiber (**Fig. 11A, C, E indicated by yellow arrow**). For example, a spiculum extended from an “octopus”-like S100B+ cell in the center of 2D-image (**Fig. 11A**) ended on TH+ varicosities. In the same image, an adrenergic neurite extended from the neuronal plexus overlapped with the spiculum of a second “octopus”-like S100B+ cell (**Fig. 11A, white star**). In **Fig 11B**, a spiculum from a S100B+ cell elongated toward a large adrenergic fiber ending with an “endfoot” dilation on the neuronal fiber. The S100B+ endings could make contacts in the shape of “endfeet” with pacemaker cells (**Fig. 11C, E**). Adrenergic and cholinergic varicosities of the neuronal plexus could also stretch alongside S100B+ processes (**Fig. 11A, B, D**). Neurites were often found in proximity (1-5µm) to S100B+ cells. Two or more neurites may approach a single S100B+ cell. Autonomic innervation did not make contact with all S100B cells in any given area, with the majority of S100B cells imaged receiving no discernible nervous input (**Fig. 11C, D**). Adrenergic and cholinergic neurites could enwrap the soma and/or spiculum of a S100B-positive cell. Cholinergic varicosities in **Figure 11** (**Panel D**) were densely present around the soma and spiculum of one S100B+ cell, which was in a cluster of 7 interstitial cells. Ameboid-like S100B+ cells were also found to be innervated (**Fig. 11C**). Finally, nerves, HCN4+ pacemaker cells, and S100B+ cells were found to make composite points of contact among each other, allowing for the possibility for communication pathways in such cellular arrangements. Extensions from multiple intertwined S100B+ cells, an HCN4+ pacemaker cell and a cholinergic neurite colocalized within 1µm of distance interacted anatomically, manifesting an apparent anatomical unit between the three cell types in **Fig 11F**. In the image, a cytoplasmic extension from an HCN4-immunoreactive cell, resembling a dendritic spine, made contact with an S100B+ ameboid pseudopodium (**white arrow in Fig. 11F**), both apparently connected varicosity from a cholinergic nerve. The **white star** in **Fig. 11F** indicates a second point of contact between the same S100B+ cells and cholinergic neurite. Finally, in the same image, the extension from a HCN4-immunoreactive cell was colocalized with a nerve ending near the nucleus of a peripheral glial cell (**indicated by white triangle Fig. 11F**). High optical zoom confocal imaging of the anatomical relationships between S100B+/GFAP- interstitial cells embedded within the HCN4+ immunoreactive pacemaker cell meshwork showed that the extensions of S100B^+^/GFAP- interstitial cells lie in close proximity to the cellular membrane of HCN4+ immunoreactive pacemaker cells, but the limited resolution of our confocal microscopy could not answer the question of the real distances between cells at the sites of these interactions. Addressing this uncertainty would shed light on the potential functions of these connections, and the types of intercellular communication in which they could be involved.

### Ultrastructure of the Interstitial Cell Populations of SAN viewed through Transmission Electron Microscopy: microenvironment and intercellular relationships

We identified interstitial glial cells within SAN semi-thin sections in TEM by the heterogenic cellular morphology and location with respect to pacemaker cells observed in our aforementioned immunohistochemical studies. These cells were located in the interstitium between pacemaker cells, had small elongated soma, and extremely long and thin cytoplasmic extensions that distinguished them from other SAN cells. These features are those of cells classified as telocytes (**Fig. 12 Panel A**), in accordance with the descriptions by Popescu and colleagues (24). The ultrastructural hallmark of telocytes is their variable cellular extensions with cytoplasmic dilations, defined as telopodes i.e., extensions, resembling “strings”, with “beads” of cytoplasmic dilations. Cytoplasmic dilations within thin fibrillar cellular extensions have been classified as podomers while the bead-like dilated cytoplasmic areas being classified as podoms (**Fig. 12 Panel B black arrowheads indicate to podoms**). Podoms accommodate organelles like caveolae, mitochondria, electron-dense granules, endoplasmic reticulum that may be involved in maintaining Ca^2+^ homeostasis. Cells with this telocyte morphology were readily identifiable in TEM images to lie within the interstitial space of the SAN (**Fig. 12 Panel B**). Further, the different types of interfaces between interstitial cells and pacemaker cells identified with immunohistochemistry (**Figures 9, 10**) could also be seen in TEM images.

**Fig. 12.**
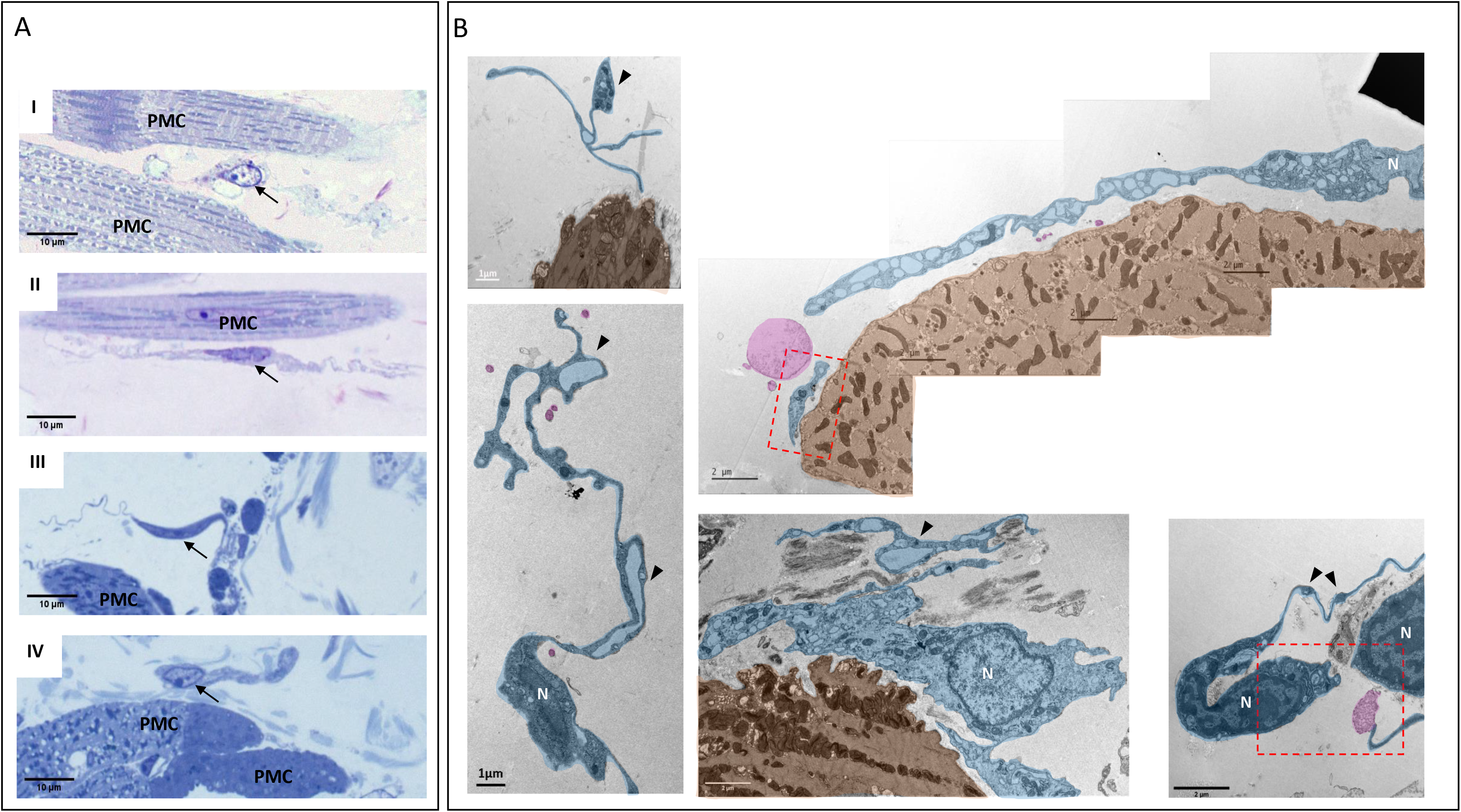
Interstitial cell populations found in EM images of thin-sections of the mouse sinoatrial node. These were identified through high resolution TEM images as telocytes, and were always found in close relationship with pacemaker cells (**PMC**). **Panel A** shows semi-thin sections of SAN stained with toluidine blue, and illustrates the heterogeneous shapes of interstitial cells, with black arrows indicating their nuclei. **Panels AI** and **AII** present bipolar cells, while piriform/fusiform unipolar cells are shown on **Panels AIII** and **AIV**. An interstitial cell with branching cellular extensions is shown in **Panel AI**. The long and thin cytoplasmic extensions of interstitial cells (telopodes) are shown in **Panels AII** and **AIII**. A flattened interstitial cell is illustrated in panel AIV. **Panel B** contains digitally colored TEM images that illustrate the ultrastructure of different types of telocytes (**blue**) located in close proximity to pacemaker cells (**brown**), as well as exosomes shed from telocytes (**pink)**. Telopode ultrastructure is characterized by extremely thin, dilated fibrillar segments (defined as podomers), and “bead”-like regions (defined as podoms, indicated by **black arrowheads**). Scarce organelles (mitochondria, endoplasmic reticulum, caveolae, and larger membranous vesicles) can be visualized in the soma and in the podoms of telocytes. Telocyte nuclei in the TEM images are indicated by the letter **N**.

Tiling of multiple electron microscopy images allowed for reconstruction of the anatomical interface between the telopodes and the surface membrane of pacemaker cells. Extracellular space between telopodes running alongside cytoplasmic membrane of a pacemaker cell varied from 0.1μm to 1μm, creating potential points of interaction. In some instances, the soma of the telocytes was located in close proximity to the cellular membrane of a pacemaker cell (**Fig. 12 Panel B lower central image**). The cellular membranes of two cells in this anatomical configuration are separated by extracellular space with smallest measured distances between membranes of 0.1μm-0.2μm **(Fig. 14)**.

**Fig. 13.**
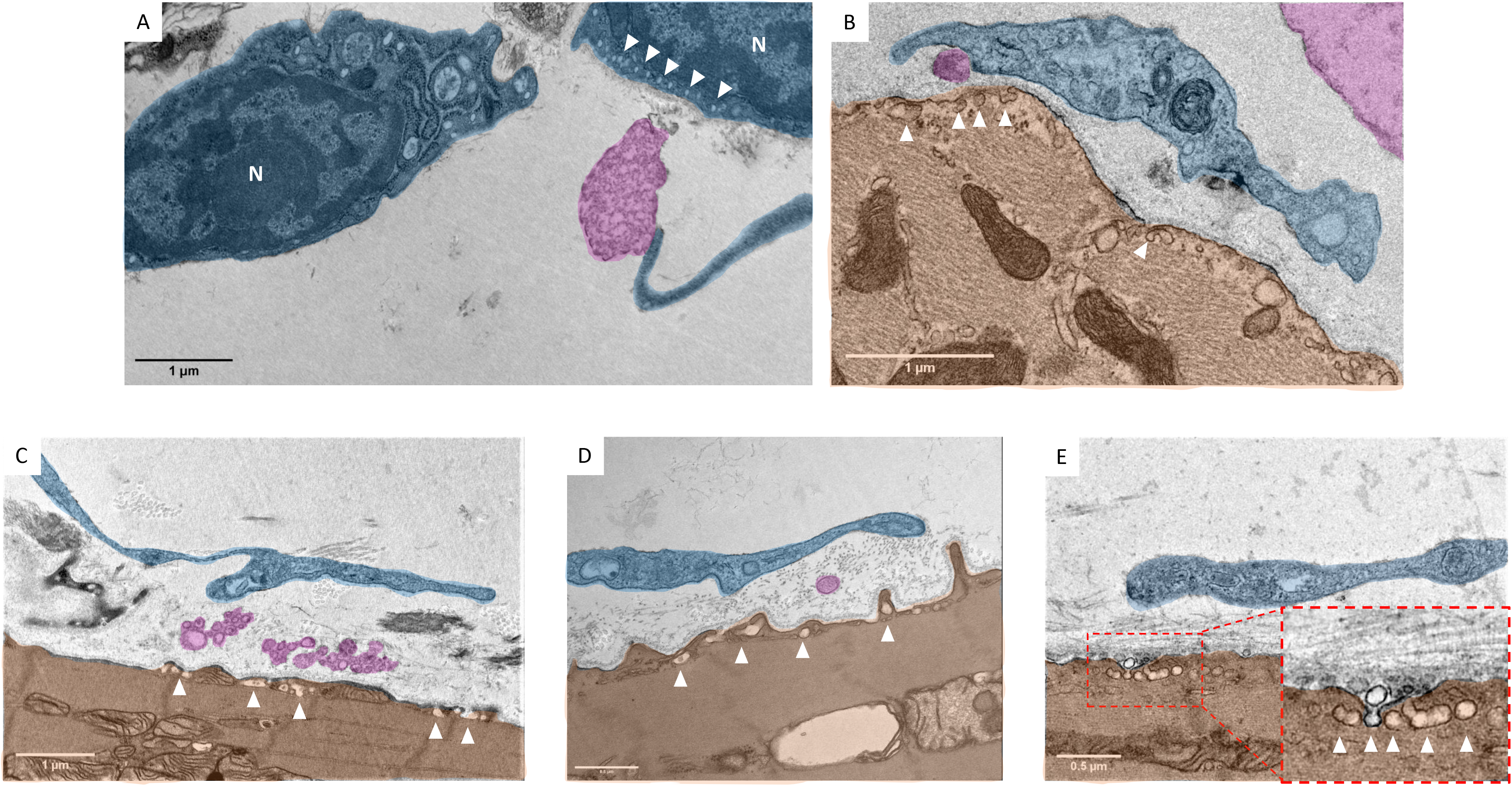
Digitally colored TEM images of “cotton” type interactions between telocytes (**blue**) and pacemaker cells (**brown**) in the SAN. **Panels A and B** illustrate zoomed-in depictions of the areas highlighted by two **red rectangles** in Figure 12 **Panel B**. **Panels C, D and E** illustrate close contacts between telopodes (**blue**) and pacemaker cells (**brown**). Exosomes and multivesicular bodies (**pink**) are easily identifiable in the intercellular space. Telocytes and telopodes could approach the plasma membrane of pacemaker cells to a vicinity of ∼200nm, but physical contact between the membranes of pacemaker cells and telocytes was never detected. Ectosomes, exosomes, and multivesicular bodies have been found in close contact to the plasma membrane of pacemaker cells. In the segments closest to telopodes, plasma membrane of pacemaker cells always had a high number of caveolae (**white arrowheads**). **Panel E** shows a virtually zoomed-in image of caveolae in the membrane of a pacemaker cell (**brown**). The nuclei of telocytes in the TEM images are indicated by the letter **N**.

**Fig. 14.**
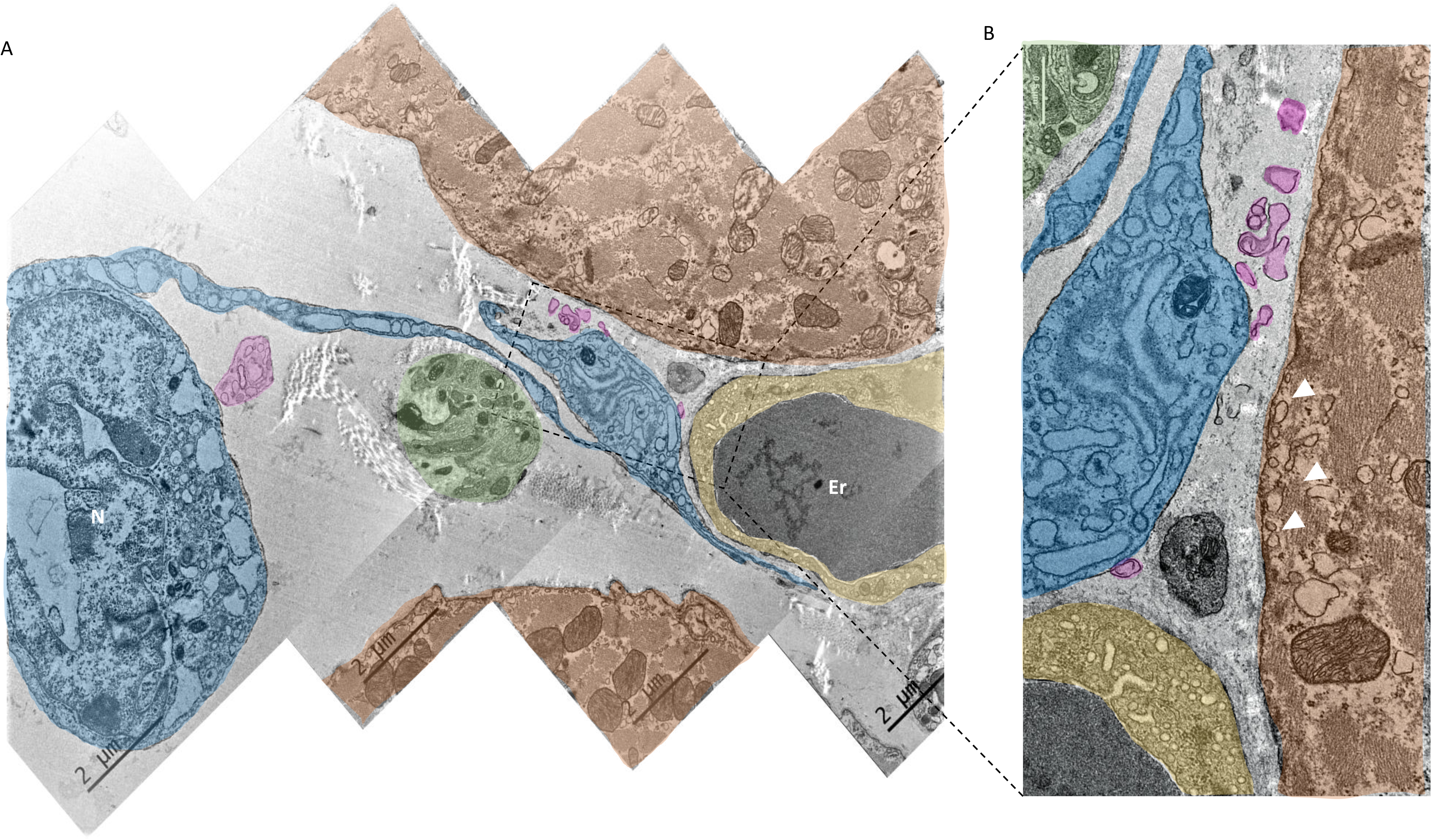
Ultrastructure of telocytes (**blue**) colocalized with neuronal fibers (**green**) and the endocardial cells of blood capillaries (**yellow**). **Panel A** illustrates a digitally colored image of SAN tiled from 4 TEM images. **Panel B** illustrates a zoomed-in picture of the area outlined by the rectangle in **Panel A**. **Panels A** and **B** illustrate close anatomical interactions between telocytes, pacemaker cells, nerves from the neuronal plexus, endocardial cells. Exosomal vesicles shed from telocytes can also be seen (**pink**). **Panels C** and **D** illustrate zoomed-in regions of interest with close contact between pacemaker cells, endocardial cells, and exosomes. A high number of caveolae (**white arrowheads**) within the plasma membrane of pacemaker cells (**brown**) were detected adjacent to a telopode (**blue**). The nuclei of telocytes in the TEM images were indicated by letter **N**. Erythrocytes in the TEM images were indicated by the letters **Er**.

Telopodes imaged by TEM showed a variable cytoplasmic diameter that could dilate and narrow across their length, a morphology shared by the S100B+/GFAP- spicula of interstitial cells seen by confocal microscopy (**Fig. 12, Fig. 13**). Telopodes were detected at distances as close as 0.2μm-1μm to pacemaker cells (**Fig. 13**). Analysis of TEM images showed the presence of ectosomes, exosomes, and multivesicular bodies (**pink** in **Fig. 12** and **13**) in the extracellular space between telopodes and the plasma membrane of pacemaker cells, strongly suggesting ongoing intercellular signaling between the two cell types. The possibility of intercellular communication between telocytes and pacemaker cells is further supported by TEM images in which highly dense plasma membrane caveolae over the surface of pacemaker cells were found in close proximity with telopodes apparently shedding vesicles towards them (**Fig. 13, caveolae are highlighted by white triangles**).

Figure 14 obtained by tiling 4 TEM images, illustrates endocardial cells of blood vessels, nerve bundles from the SAN neuronal plexus, telocytes, and pacemaker cells colocalized within an area of 15μm by 20μm in which telopodes could be found approaching nearby cells at distances as close as ∼200nm (**Fig. 14**). Zoomed-in TEM images show exosomes apparently shed by telocytes toward both endocardial and pacemaker cells (**Fig. 14 Panel C**).

### Effect of S100B on action potential induced Ca^2+^transients generated within HCN4+ pacemaker cells

The intertwining of S100B+/GFAP- interstitial cells and pacemaker cells within the HCN4+ meshwork interstitium, the congruent cellular phenotypes of S100B+/GFAP-interstitial cells and telocytes seen in EM and apparent vesicular secretion from telocytes towards pacemaker cells seen in EM images suggest S100B may influence pacemaker function. We had previously established that local calcium releases (LCRs) and action potential induced calcium transients (APCTs) within and among HCN4+ cells were critical events underlying pacemaker function (9). Given that Ca^2+^ dynamics are crucial to SAN impulse generation, and that S100B is a secreted EF-Hand protein with potent Ca^2+^ buffering capacity and signaling properties (25) via which glial cells communicate to CNS neurons to alter their firing patterns (26, 27), we hypothesized that S100B applied to the SAN would alter Ca^2+^ dynamics in tissue, and therefore impact the SAN APCT firing rate and rhythm. To test the hypothesis of a functional relationship between S100B+/GFAP- interstitial cells and HCN4^+^ pacemaker cells we measured the effect of applied S100B protein on LCRs and APCTs in SAN tissue. We used HCN4-GCaMP8 transgenic mice, which express a genetically encoded Ca^2+^indicator exclusively within HCN4+ pacemaker cells to image Ca^2+^dynamics in ex-vivo tissue preparations with high spatial and temporal resolution.

The earliest APCTs in the SAN shown in **Figure 15** (**Panel A**) and **Supplemental Movie 1** were detected near the crista terminalis within the “body” of the HCN4+ meshwork, closer to the SVC (**right**), than to the IVC (**left**). “Chronopix” analysis (which color-coded the phase shift between APCTs in different time areas, as in (9)) showed that HCN4+ cells with different phase shifts relative to moment of occurrence were intertwined: following the earlier APCT occurrence, subsequent APCTs were generated toward the SVC and IVC simultaneously (**Panel C**).

**Fig. 15.**
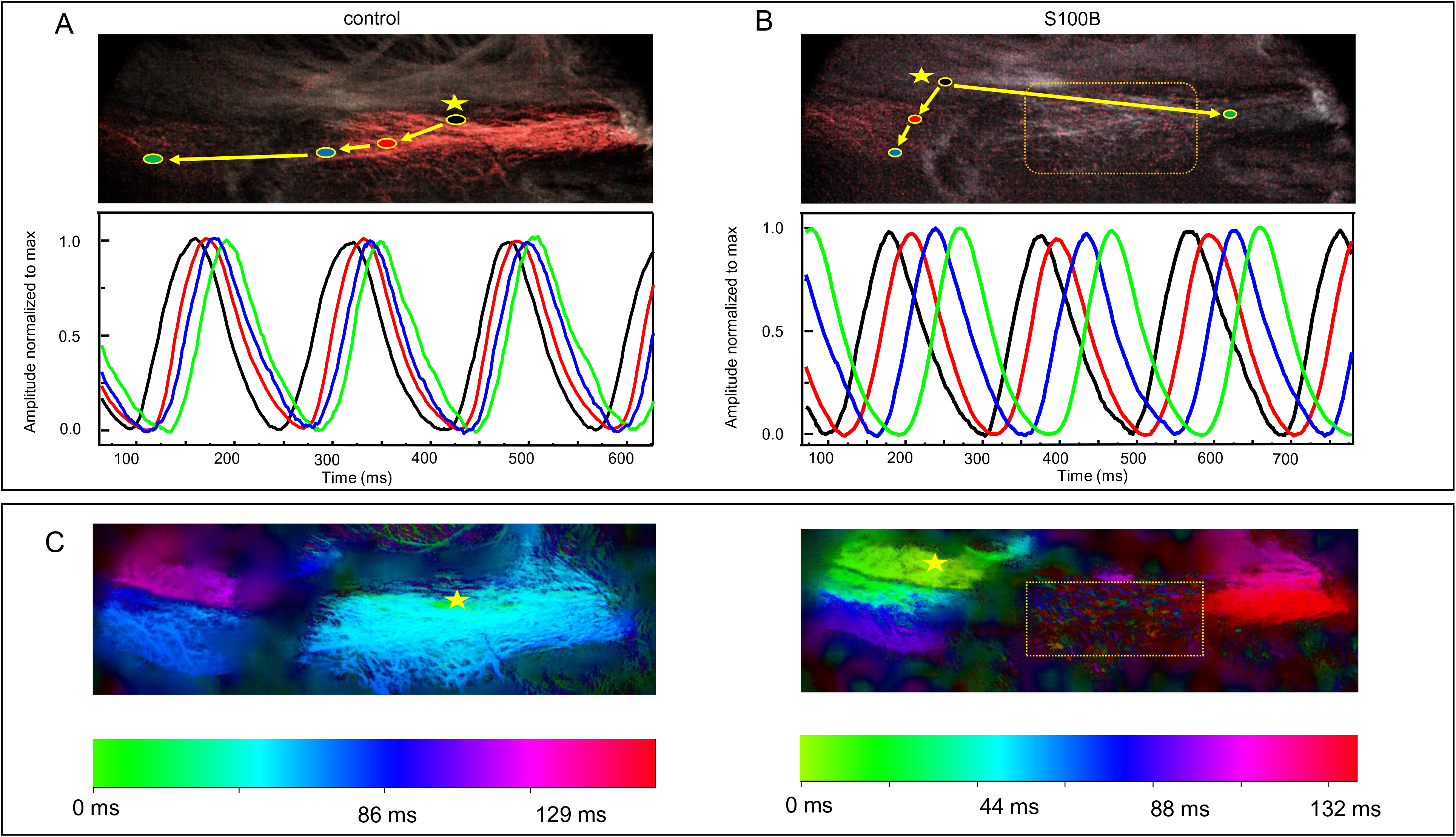
2D-images of action potential induced calcium transients (APCTs) detected within pacemaker cells expressing the genetically encoded Ca^2+^ indicator GCaMP-8. The region of SAN highlighted by a **yellow rectangle** stopped generating APCTs and only exhibited local calcium releases (LCRs) after the addition of S100B. The images shown in **Panels A** and **B** depict the instances of highest fluorescence from recordings made under control conditions, and 20 mins after the addition of 200nM S100B. The increase in fluorescence of genetically encoded Ca^2+^ indicator is highlighted by **red** color. The site of detection of the first APCTs is marked by a **yellow star** in all corresponding 2D- Ca^2+^ images. The direction of the consequent appearance of APCTs is indicated by **arrows**. In the fluorescence tracings under each image, color coding of the APCTs corresponds to the color of the region of interest depicted on the 2D-calcium image above the superimposed APCTs plot. The **black line** corresponds to the first detectable APCT, and the area of the SAN with the corresponding first APCT is marked by a **black ROI** within a **yellow circle** in the 2D-calcium image. **Panel C** illustrates chronopix analysis performed on the same recording as the previous panels.

200nM S100B added to the superfusate shifted the APCT initiation site toward the IVC (n=4, **Fig 15 Panel B**). Moreover, following the addition of S100B to the superfusate, areas in which the earliest APCTs were observed in control became devoid of APCTs, and in this area only LCRs were detectable (**Fig. 15 Panels B, C; yellow square**). The time delay between the earliest APCTs (**Fig.15 A, B**) and those detected at the root of the IVC, measured by the phase shift at the point of halfwidth between the APCTs, averaged 67.2+10.4ms (n=4), and the APCT firing rate averaged 405+19 beat/min, in control conditions. S100B slowed the average APCTs firing rate to 307±30 beats/min after 31.7±5.4 minutes. Removal of S100B from the perfusate increased the APCT firing interval to 322±34 beats/min.

### S100B desynchronizes SAN Ca2+ signaling, increases the cycle length variability, and prolongs the mean cycle length

In order to determine the synchronicity among APCTs generated by HCN4+ cells we measured APCTs at low-zoom, which summates the ensemble of APCTs generated within HCN4+ across the entire SAN during each cycle, informing on mean APCT firing interval, the APCT interval variability, and the kinetics of APCT formation and decay during each cycle. In other terms, each low-zoom APCT reports an integrated version of the chronopix shown in **Fig 15 Panel C**, during the formation and decay of APCTs. Together, the chronopix map and the low-zoom APCT tracings in **Figure 16** provide some idea of **2D spatial-temporal** integration of events occurring during each APCT cycle. The time to peak amplitude, the maximum rate of rise measured at low zoom inform on synchronicity of development of APCTs across the entire SAN, and the maximum decay rate informs on the synchronicity of APCT decay rates. Given its Ca2+ buffering capacity we hypothesized that S100B would prolong the mean cycle length and increase the APCT cycle length variability, while reducing synchronicity of APCT development and decay during each APCT cycle (28).

**Fig. 16.**
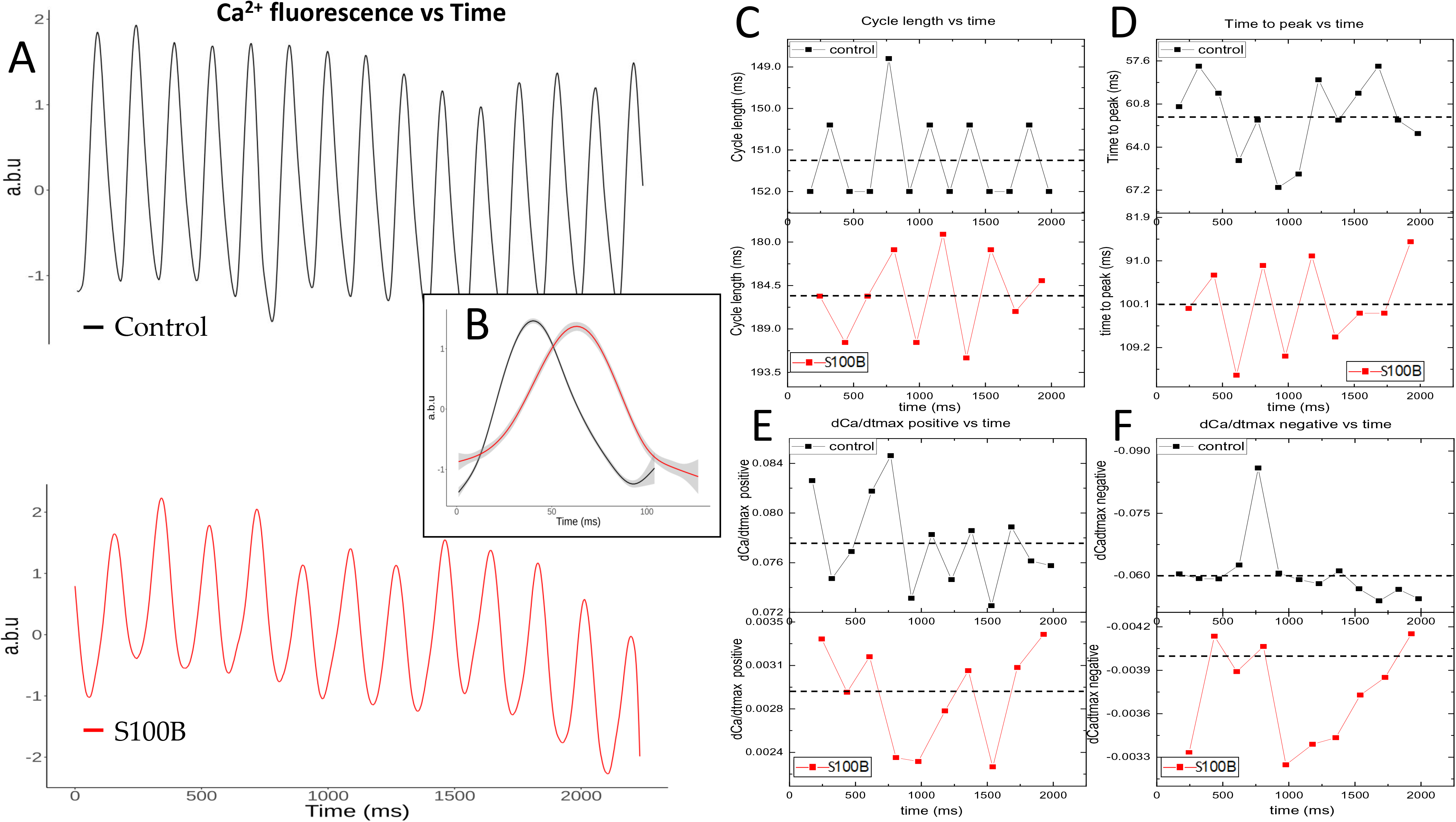
APCT tracing analysis from a representative recording of Ca^2+^ transients, obtained by plotting the intensity of fluorescence (measured in arbitrary intensity units, a.b.u.) detected within pacemaker cells expressing GCaMP-8 before and after addition of S100B to the superfusion (n=3). **Panel A** depicts normalized APCTs measured at low-zoom, in control conditions (**black line**) and 20 minutes after addition of 200nM S100B (**red line**). A **dashed line** indicates the mean value for every parameter. **Panel B** represents calculated average normalized APCT waveforms for both the tracings, superimposed. **Gray shading** displays the standard error along the length of the APCT. **Panel C** shows the variability APCT cycle lengths and the mean APCT cycle length for all APCTs occurring in the time series. The mean cycle length increased from 151.3ms to 185.6ms after addition of S100B, with its range increasing from 4ms. to 12.8ms. **Panel D** shows the time from APCT initiation to peak amplitude, plotted over time for both conditions in. The time to peak increased from 61.8 ms to 100.1 ms after addition of S100B, with its range increasing from 9ms to 28ms. **Panel E** shows the maximum rate of rise from APCT initiation to peak, or dCa^2+^/dt_max_ positive, plotted over time for both conditions. The mean maximum rate of rise decreased from 0.078a.b.u./s to 0.0029a.b.u./s after addition of S100B, with its range decreasing from 0.01a.b.u./s to 0.001a.b.u./s. **Panel F** shows the maximum rate of decay from APCT peak to the end of the cycle, or dCa^2+^/dt_max_ negative, plotted over time for both conditions. The mean maximum rate of decay decreased from 0.061a.b.u./ms to 0.0037a.b.u./ms after addition of S100B, with its range decreasing from 0.042a.b.u./s to 0.001a.b.u./s.

**Figure 16** shows the impact of S100B on these APCT characteristics in the same representative SAN preparation shown in **Fig 15**. **Supplemental Tables 1-3** list the effects of S100B on these APCT characteristics in the 3 SAN preparations to which S100B was applied. **Figure 16 Panel A** shows a time series of APCTs prior to and during S100B superfusion. **Panel B,** which depicts superimposed normalized average APCT waveforms for the time series, illustrates the marked impact of S100B on the shape of APCTs, including prolongation of the time to peak amplitude and a marked reduction in the maximum rate at which low zoom APCTs developed, and a marked reduction in the maximum rate of APCT decay. **Panel C** demonstrates that S100B not only increased the cycle to cycle variability of APCTs within the time series, but also markedly prolonged the mean APCT interval across the time series. Note that the degree of regularity of the APCT cycle length variability seen in control is distorted, and virtually absent, in the presence of S100B. S100B increased the time from APCT initiation to peak amplitude in all three tested preparations (**Panel D**), and significantly decreased both the maximum rate of rise **(Panel E**) and the maximum rate of decay (**Panel F**) of APCTs. These effects on cycle length, time to peak, maximum rate of rise, and maximum rate of decay were consistent and significant in all tested preparations (**Supplemental Tables 1-3**). Further, S100B markedly increased the range cycle length and time from APCT initiation to peak amplitude across the time series (**Fig 16 Panels C and D, Supplemental Tables 1-3**).

The types of cells and cellular interactions identified in our studies are summarized in **Table 1**.

## Discussion

Enabled by coupling 3D confocal tile imaging techniques with various combinations of triple immunolabeling for S100B, GFAP, HCN4, TH, and ChAT (or VAChT), to optically dissect the mouse SAN, we reconstructed 3D images of the entire SAN from 3D optical sections, IVC to SVC, endocardium to epicardium, at multiple levels of magnification. Thus, we obtained both panoramic and finely detailed images of local SAN cytoarchitecture that describe not only the distribution of different cell types throughout SAN tissue, but also their variable phenotypic details and connectivity patterns among each other.

We had previously noted that the HCN4+/CX43- meshwork is characterized by cells with multiple branches and extensions of variable size and shape, resembling the arborization of brain neurons (**Table 1**) (9). Further, we described the presence of an HCN4-/CX43+ network within the SAN, that locally interacted with cells in the HCN4+ meshwork. We also noted that multiscale, local heterogeneous Ca^2+^ signals within and among these cells underlie the generation of impulses that exit the SAN to initiate each heartbeat. This complex cytoarchitecture of SAN tissue and heterogeneous local Ca^2+^ signaling patterns within and among SAN pacemaker cells strikingly resemble those of brain neuronal networks that generate nerve impulses (9).

Here, we uncovered novel aspects of the autonomic plexus within the SAN, and discovered the existence of two novel structures that intertwined with the HCN4+ meshwork: 1) a web of S100B+/GFAP+ peripheral glial cells, and 2) a network of S100B+/GFAP- interstitial cells. Combining previous research and the work presented here, the SAN cytoarchitecture may be defined as having at least 5 interacting 3D cell networks, composed of interacting cell types: HCN4+/CX43- pacemaker cells, HCN4-/CX43+/F-actin+ cells (9), TH+ or VChAT+ autonomic nerves, GFAP+/S100B+ peripheral glial cells, and S100B+/GFAP- interstitial cells (See **Table**). This cytoarchitecture suggested to us that these numerous networks may communicate with each other to influence the intra-SAN pattern of Ca^2+^ dynamics that underlies AP firing rate and rhythm.

### ICG and Neuronal Plexus Communication with the SAN Cytoarchitecture

Although nerves extending from ICG neurons are known to be essential participants in the communication between the autonomic nervous system and SAN cells, our novel findings regarding the cytoarchitecture of the SAN imply more complex signaling than heretofore appreciated, beyond that of neurites to pacemaker cells. Images of ICG stained for TH, VAChT, and HCN4, confirmed that most neurons in these ganglia were cholinergic (i.e., immunoreactive for VAChT), while others were adrenergic, and that some neurons within ICG were reactive for neither VAChT nor TH (**Fig. 2**), suggesting the presence of other neurotransmitters within ganglionic neurons (18). We also observed HCN4+ immunoreactivity on the shell of ICG suggesting that some HCN4+ satellite cells may be present around the ganglia. HCN4+ pacemaker cells were not detected within SAN around the ICG (**Fig. 2**).

The results of our panoramic optical mapping of the autonomic plexus in the HCN4+ meshwork provide novel perspectives on the innervation originating in ICG neurons to penetrate the SAN interior (**Fig. 3**). Although the number of nerve fibers innervating the SAN is high throughout the tissue, the distribution of autonomic neurites showed substantial heterogeneity, suggesting that HCN4+ cells within different areas of the SAN likely vary in excitatory or inhibitory autonomic input, that would likely provide differential local control of pacemaker cell beating rate and ultimately impact the emergent properties of SAN function as a whole. These adrenergic and cholinergic innervation patterns likely create areas of dominant excitatory innervation, in which the local AP firing frequency is regulated by adrenergic input in the absence of cholinergic inhibition, while in adjacent parts of the meshwork adrenergic input is likely balanced by acetylcholine release from cholinergic synapses. Because sympathetic nerves may innervate the heart bypassing the ICG, areas of dominant adrenergic input imply the possibility of nervous signals reaching the SAN without modulation by the ICG (29). The presence of long-recognized accentuated antagonism (30) attributable to vagal stimulation relative to sympathetic stimulation likely results not only from the dominance of cholinergic input in the ICG, but also the local variability of autonomic innervation. Because cells of the HCN4+ meshwork are defined by a constitutively active adenylate cyclase which impacts on intracellular Ca^2+^ dynamics to favor a faster beating rate, the net inhibitory effect of neuronal input into the SAN counterbalances this intrinsic *modus operandi* of pacemaker cells. The heterogeneity in autonomic innervation of the SAN could also be involved in the shifts in the site of earliest impulse initiation within the SAN observed previously in response to application of autonomic neurotransmitters or selective stimulation of vagal or sympathetic fibers. Differential autonomic input also likely impacts differences in the microvascular pattern observed among different SAN areas (31).

### Communication between the Novel Peripheral Glial Cell Web and Novel S100B+/GFAP- Interstitial Cell Network with the SAN Cytoarchitecture

We were on a quest to find CNS-like neuron-glial cell coupling interactions within the SAN that extend **beyond** neurites of nerve fibers that emanate from neuronal ganglia embedded within the SAN epicardium. The S100B+/GFAP+ peripheral glial cells (PGCs) that traversed the SAN’s “head”, “body” and “tail” were arranged in a regular web pattern across the entire HCN4+ meshwork, which has never been previously visualized (**Fig. 6**). We also found the somata and projections of some PGCs to be closely associated with the endothelial cells outlining the lumen of blood vessels within SAN tissue, which hinted at the existence of a peri-vascular interface within the heart similar to the perivascular interface within the brain. This discovery hinted to us the possibility of integrated regulation of blood vessel diameter, and therefore blood flow, together with pacemaker cell firing and SAN energy consumption.

Further, when searching for GFAP+/S100B+ PGCs, we also discovered a remarkable, novel, population of S100B+/GFAP- cells that was irregularly distributed, some individually and in some in groups or clusters across the HCN4+ meshwork, (**Fig. 7**). S100B+/GFAP- cells were characterized by somata of variable size (∼10µm in diameter), and strikingly diverse cellular morphology, suggesting that this expression phenotype represents multiple cell types (**Fig. 8, 9**), thus indicating a novel component of SAN cytoarchitecture. Multicellular complexes of S100B+/GFAP- cells, HCN4+ pacemaker cells, the other novel GFAP+/S100B+ PGC network described here, and neurites emanating from autonomic neurons create anatomical, and likely functional, units within the SAN.

Employing TEM to define the types of connections between pacemaker cells and other cells of the node more precisely, we observed interstitial cells with the distinct morphology of telocytes, resembling in many ways the long projections of S100B+/GFAP- interstitial cells. TEM images also demonstrated the presence of interstitial exosomes apparently shed from telopodes (**Fig. 13 and Fig. 14**), that were in close vicinity (0.2-0.5µm) to caveolae on the surface membrane of pacemaker cells, hinting at vesicular release and uptake in the synaptic structures between telocytes and HCN4+ cells. These discoveries strongly suggested to us that telocytes may play a central role in establishing a multicellular complex within the SAN by providing both anatomical and functional coupling between different SAN cell types. In the absence of immunolabeling of TEMs, however, the certainty that these interstitial telocytes are the very same interstitial cells identified to express S100B in our triple immunolabeled 3D confocal images cannot be verified and will require additional studies using immunogold labeling.

### A Role of S100B in Intercellular Communication within the SAN Cytoarchitecture

The intertwining of S100B expressing cells and pacemaker cells within the HCN4+ meshwork, and the presence of vesicles within telocytes (**Fig. 14, 15**), suggesting apparent vesicular secretion from telocytes into the interstitial space between them and HCN4+ pacemaker cells, makes it likely that telocytes communicate with HCN4+ pacemaker cells via exosomal secretion.

While S100B is typically expressed in the glial cells, it has also been detected in other cell types, all of which originate from neural crest cells, e.g., melanocytes (32) and adipocytes (33). As a member of the large S100 protein family, S100B possess two EF-hand Ca^2+^ binding motifs, conveying them the ability to buffer Ca^2+^ (25), which may be implicated in its effect to markedly alter Ca^2+^ signaling and impulse generation and its rhythm that we observed in SAN preparations (**Figs. 16, 17**). Thus, communication between S100B-expressing cells and pacemaker cells within the SAN to alter intracellular Ca^2+^ signaling may be implicated in the initiation of the heartbeat. S100B secreted from glial cells has been implicated in the modulation of rhythmic activity in trigeminal sensory-motor circuit for mastication (26) as well as in enhancement of kainate-induced gamma rhythmogenesis in hippocampal CA1 (27), highlighting the crucial importance of glial cell secreted proteins and gliotransmitters min modulating the AP firing rate and rhythm within neural circuits (34). Further, S100B contributes to nerve sprouting and can be used to assess the health of the intrinsic cardiac autonomic nervous system (35). In this regard, a recent study of the catheter-based treatment of atrial fibrillation established that release of S100B from cardiac glia is a hallmark (35) of neural damage.

We had previously established that self-organization of LCRs in single SAN pacemaker cells leads to the generation of action potentials (36), and that LCRs and APCTs within and among HCN4+ cells in SAN tissue were critical events determining pacemaker function. Because S100B expressing cells and HCN4+ pacemaker cells are intertwined within the SAN cytoarchitecture, we reasoned that secretion of S100B, a potent Ca^2+^ buffering protein, may influence SAN pacemaker function. Indeed, addition of 200nM S100B to the perfusate of ex-vivo whole mount SAN preparations had a marked effect on the signaling of HCN4+ cells, which was associated with changes in the site of earliest AP appearance within the SAN (i.e., shifting from the “body” of the SAN to the “tail”). S100B also modified the local kinetics and patterns of subsequent APCT occurrence in other regions of the SAN, increasing the variability in beat-to-beat calcium kinetics and slowing down the onset of each APCT. In fact, addition of S100B buffered Ca2+ signal transduction to such an extent that the central body of the SAN began to lack coordinated APCTs, and only exhibited LCRs (**Fig 15, Supplemental Figure 1**).

The degree of synchronization among molecular functions that regulate the coupled-clock system intrinsic to pacemaker cells is manifest, in part, in the self-ordering (synchronization) of LCRs that occurs during the evolution of the electrochemical signal that is otherwise known as spontaneous diastolic depolarization of SAN cells in isolation. The kinetics of the formation and decay of coupled voltage and Ca^2+^ signals during a given AP cycle, the AP firing interval variability, and the mean number of action potentials fired per unit time are self-similar to each other (28). We envisioned that in SAN tissue, the appearance of an APCT within each chronopix (**Fig 15 Panel C**) informs on the emergence and continuation of spatial-temporal synchronization of Ca^2+^ signals during a given beat, speculating that APCT development in SAN tissue resembles the self-ordering of LCRs seen in isolated single cells during AP cycles (37).

The present results demonstrate that S100B applied to SAN tissue not only impacted on the spatial-temporal distribution of emergent APCTs across the SAN during each impulse formation, as seen in chronopix maps, but also increased the inter-impulse variability (**Fig 16 Panels A, C, and D, Supplemental Tables 1-3**). Further, S100B markedly altered the kinetics within each low-zoom APCT, markedly prolonging the time from impulse initiation to its peak amplitude and markedly reducing the maximum rates of rise and decay (**Fig 16 Panels A, C, D, E, F Supplemental Tables 1-3**).

Because intracellular Ca^2+^ levels in pacemaker cells are highly dependent on Ca2+ flux into and out of the cell during each beat, we cannot ascertain from our results whether these marked effects of S100B on synchronicity of SAN APCTs and beat to beat APCT cycle length variability and mean cycle length, result from intracellular or extracellular Ca^2+^ buffering, or a combination of both. Future experiments are required to determine the mechanism by which S100B affects Ca^2+^ dynamics in the SAN whether extracellular or intracellular, or even whether it is exclusively limited to its buffering properties.

### Conceptualizing the SAN as the heart’s “Central Brain”

Our discoveries reported here, together with those in our prior study (9), provide the basis for describing the cytoarchitecture and function of SAN tissue as distinctly neuronal-like. The variability and complexity of cellular phenotypes and intracellular calcium signaling kinetics found in the SAN more closely resemble the cytoarchitecture and calcium signals of neuronal tissue: pacemaker cells are connected within the meshwork with branching cellular extensions, similar to the dendritic arborization of neurons; the calcium signaling found in the SAN, with its variable amplitudes, frequencies, kinetics, is also conceptually similar to the mechanisms controlling AP-mediated signaling observed in neuronal tissue.

The heterogeneous pattern of pacemaker cell interconnectivity combined with the variable scattering of S100B+ interstitial cells (see **Table**) may modulate local Ca^2+^ signal transduction throughout the HCN4+ meshwork. Our experiments support this hypothesis, demonstrating that S100B applied to ex-vivo SANs reorganized the initiation and formation of the SAN impulse, suggestive of the type of modulatory interactions observed between astrocytes and neurons.

Thus, we conceptualize pacemaker cells within the HCN4+/CX43- meshwork as **“neuromimetic”**, sharing phenotypical similarities with neurons within brain neuronal networks. Therefore, in order to conceptualize the SAN (here depicted as intertwined networks of autonomic neurites, HCN4+-pacemaker cells, peripheral glial cells, S100B+ interstitial cells, and telocytes) as the heart’s “**central brain,**” which is not to be confused with the “little brain of the heart” (2), known as **ICG,** that receives inputs from the autonomic nervous system and which modulates the rate of AP firing of pacemaker cells via release of transmitters from synaptic terminals within the SAN neuronal plexus. In conceptualizing the SAN as the heart’s “**central brain**”, we will borrow terms utilized to describe cells and structures within nervous tissue, even though the fine details of these terms have not yet been fully defined for the SAN. Rather, the heart’s **“central brain**” integrates signals not only from the ICG, but also from peripheral glial cells and local mechanical factors within the SAN. It then forms a near-term memory of these factors through the same mechanisms observed in isolated pacemaker cells i.e., post-translational modifications, due largely to Ca^2+^ dependent phosphorylation of coupled-clock proteins that regulate their AP firing rate and rhythm (28, 38). This near-term memory, formed over the time frame of several beats, instructs the SAN on when to initiate the heartbeat (28).

Heterogeneous colocalization of different cell types within the anatomical units defined above enables the exchange of signaling molecules that drive and transform local and global calcium signals. Similar to neuronal networks (39, 40), the HCN4+ meshwork both generates and post-processes Ca^2+^ signals. Thus, we envision cardiac rhythmogenesis to result from brain-like integrative structure and function having near-term memory, rather than from the dominant activity of pacemaker cells with highest frequency of spontaneous APs.

The “neuromimetic” meshwork of HCN4+ pacemaker cells does not constitute the entire SAN. We had previously described a network of HCN4-/CX43^+^/F-actin^+^ striated cells, which (9) intertwines with the HCN4^+^ meshwork. These striated cells, therefore, contain elements that relate to muscle contractility, as well as “neuromimetic” spontaneous calcium activity. Points of contact between HCN4+/CX43- cells and striated HCN4-/CX43+/F-actin+ may establish a conversion from the apparently conducted heterogeneous electrochemical signals generated within the HCN4+ meshwork to a more ordered form of electrical conduction within striated cells, as noted previously (9). It is also likely that feedback signaling from HCN4-/CX43+/F-actin+ striated cells to the HCN4+ meshwork impacts the consequent self-ordering of its electrochemical signals. In other terms, each heartbeat is initiated by heterogeneous, local Ca^2+^ oscillations within and among numerous types of SAN cells, including striated F-Actin+/CX43+ cells, HCN4+/CX43- cells, and likely the glial cell web. The “neuromimetic” calcium signaling within and among the cell types in these cell meshes/nets/webs becomes self-organized, by mechanisms yet to be understood, to produce the net impulse activity that exits the SAN towards cardiomyocytes. The fact that Ca^2+^ signals are informed and transformed by such a variety of cell types grouped into anatomical units strengthens our vision of the SAN as the heart’s **“central brain,”** makes for a strong hypothetical framework for future research on the sinoatrial node.

THE END

## Supplement

### Methods

#### Whole mount SAN Immunolabeling

We employed extensive immunolabeling of whole-mount mouse SAN tissue preparations to reveal the complex cytoarchitecture of the SAN. A subset (n=7) of SAN preparations was fixed in 4% paraformaldehyde overnight at 4°C. The SANs were washed three times in phosphate-buffered saline (PBS) and permeabilized overnight in PBS containing 0.2% Triton X-100 and 20% DMSO, followed by additional 24 hrs permeabilization in PBS containing 0.1% Triton X-100, 0.1% Tween 20, 0.1% sodium deoxycholate, 0.1% NP-40 and 10% DMSO. After blocking the non-specific binding sites by incubation for 24 hrs. With 0.2% Tween-20 in PBS containing 3% normal donkey serum (NDS), SAN whole-mount preparations were incubated for 3 days with the primary antibodies diluted in PBS containing 0.2% Tween-20 and 3% NDS. They were then washed three times with PBS containing 0.2% Tween-20, incubated overnight with appropriate secondary antibodies, and washed three times with PBS containing 0.2% Tween-20. Whole-mount SAN preparations were mounted in Vectashield (Vector Laboratories, Burlingame, California) and sealed with a coverslip. The SAN preparations were mounted with the endocardium uppermost. Immunolabeling of whole-mount SANs was imaged with a Zeiss LSM800 confocal fluorescence microscope equipped with Plan-Apochromat 20x/1.0 Corr DIC M27 or Plan-Apochromat 40x/1.0 DIC M27 water immersion objectives and appropriate lasers and filters for fluorescence imaging (Carl Zeiss, Oberkochen, Germany). To obtain high resolution images of the whole SAN, the preparations were imaged using the tile scanning mode which acquires images (tiles) that are adjacent in x and y direction to each other without an overlap. Each tile region was based on an X and Y coordinate of the stage, and for 3D visualization, on a Z coordinate of the focus drive. Typically, optical sectioning (Z stacking) at 1 µm Z-steps from to a depth ∼100 µm of SAN tissue was used to capture individual tiles into a Z-stack.

#### Electron Microscopy Methods

Sinoatrial nodes were processed for transmission electron microscopy visualization. Fixation for electron microscopy was performed using 2.5% glutaraldehyde in 0.1 M sodium cacodylate buffer, pH 7. Samples were post fixed in 1% osmium tetroxide for 1 h at 4°C in the same buffer, dehydrated and then embedded in Embed 812 resin (Electron Microscopy Sciences, Hatfield, PA) through a series of resin resin-propylene oxide gradients to pure resin. Blocks were formed in fresh resin contained in silicon molds, and the resin was polymerized for 48-72 h at 65°C. Blocks were trimmed and sectioned in an EM UC7 ultramicrotome (Leica Microsystems, Buffalo Grove, IL) to obtain both semi-thick (0.5-1 µm width) and ultrathin (40-60 nm width) sections. Semi-thick sections were mounted on glass slides and stained with 1% toluidine blue in a 1% borax aqueous solution for 2 min. Micrographs were obtained using a Leica AXIO Imager light microscope with a Axiocam 512 color camera (Carl Zeiss, White Plains, NY). Ultrathin sections were stained with uranyl acetate and lead citrate, and then imaged on a FEI Tecnai G^2^ 12 Transmission Electron Microscope (TEM) with a Gatan OneView 16 Megapixel Camera.

#### APCT Analysis for S100B superfusion experiment

The intracellular Ca^2+^ signals recorded from whole SANs were transferred to 2D recordings of Ca^2+^ fluorescence over time and AP-induced Ca^2+^ transient waveforms were produced for further analysis. Selected AP-induced Ca^2+^ transient parameters were measured then by pCLAMP 10.4 (Clampfit) software and via in home customized program (41).

AP-induced Ca^2+^ transient cycle length was measured as the time interval between the peaks of two adjacent AP-triggered Ca^2+^ transients. Other AP-triggered Ca^2+^ transient’s parameters included the Ca^2+^ transient change over time as positive and negative dCa/dtmax (F(/F_0_*s)). Data are displayed as mean, SD and range (min and max value). Statistical significance between control and 100B was tested by Student’s *t-*test. P<0.05 was considered statistically significant. The individual normalized AP-triggered Ca^2+^ transients were isolated and averaged into a single representative wave using a smoothed conditional mean function (geom_smooth in ggplot2, R version 3.6.1) to superimpose the S100B and Control waveforms.

**Supplemental Figure 1:**
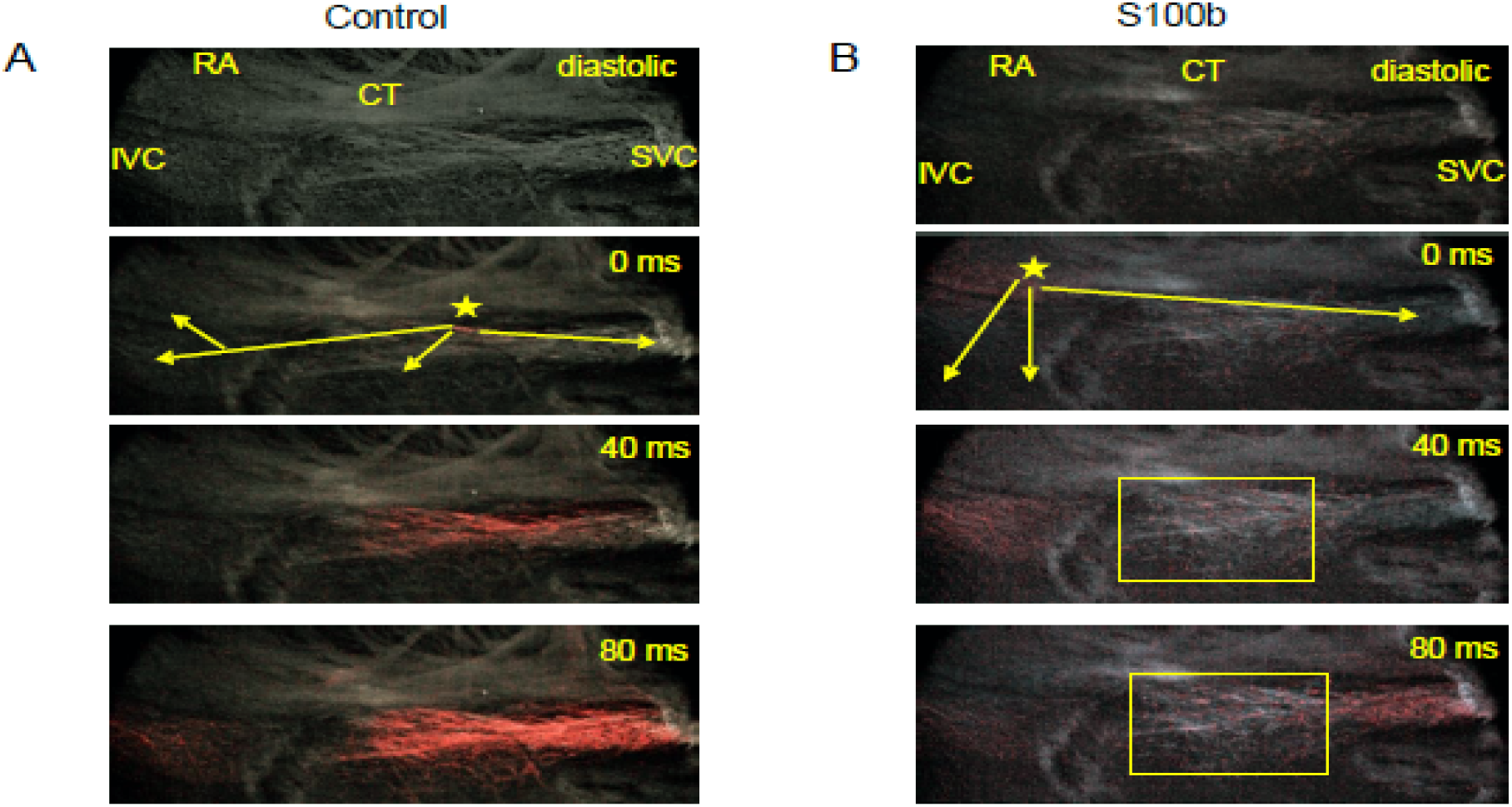
Propagation of APCTs during an S100B superfusion experiment, in a sample recording.

**Supplemental Table 1:** Experiment 1 Mean, SD and range of Ca^2+^ transient parameters in control and during incubation with S100B. Experiment 1: Ca^2+^ transient parameters in control and during incubation with S100B (mean, SD and range).

|  | Control (13 cycles) |  |  | S100B (10 cycles) |  |  |
| --- | --- | --- | --- | --- | --- | --- |
|  | Mean | SD | Range | Mean | SD | Range |
| Cycle length (ms) | 151.30 | 1.06 | 148.0-152.0 | 185.60*** | 4.46 | 179-192*** |
| Time to peak (ms) | 61.77 | 2.89 | 58-67 | 100.10*** | 9.27 | 87-115*** |
| dCadtmax positive ($F/(F_0*s)$) | 0.0776 | 0.0037 | 0.07-0.08 | 0.0029*** | 0.0004 | 0.002-0.003*** |
| dCadtmax negative ($F/(F_0*s)$) | 0.0611 | 0.0081 | 0.05-0.008 | 0.0037*** | 0.0004 | 0.003-0.004*** |

\*\*\* $p < 0.001$ , S100B vs control.

**Supplemental Table 2.**
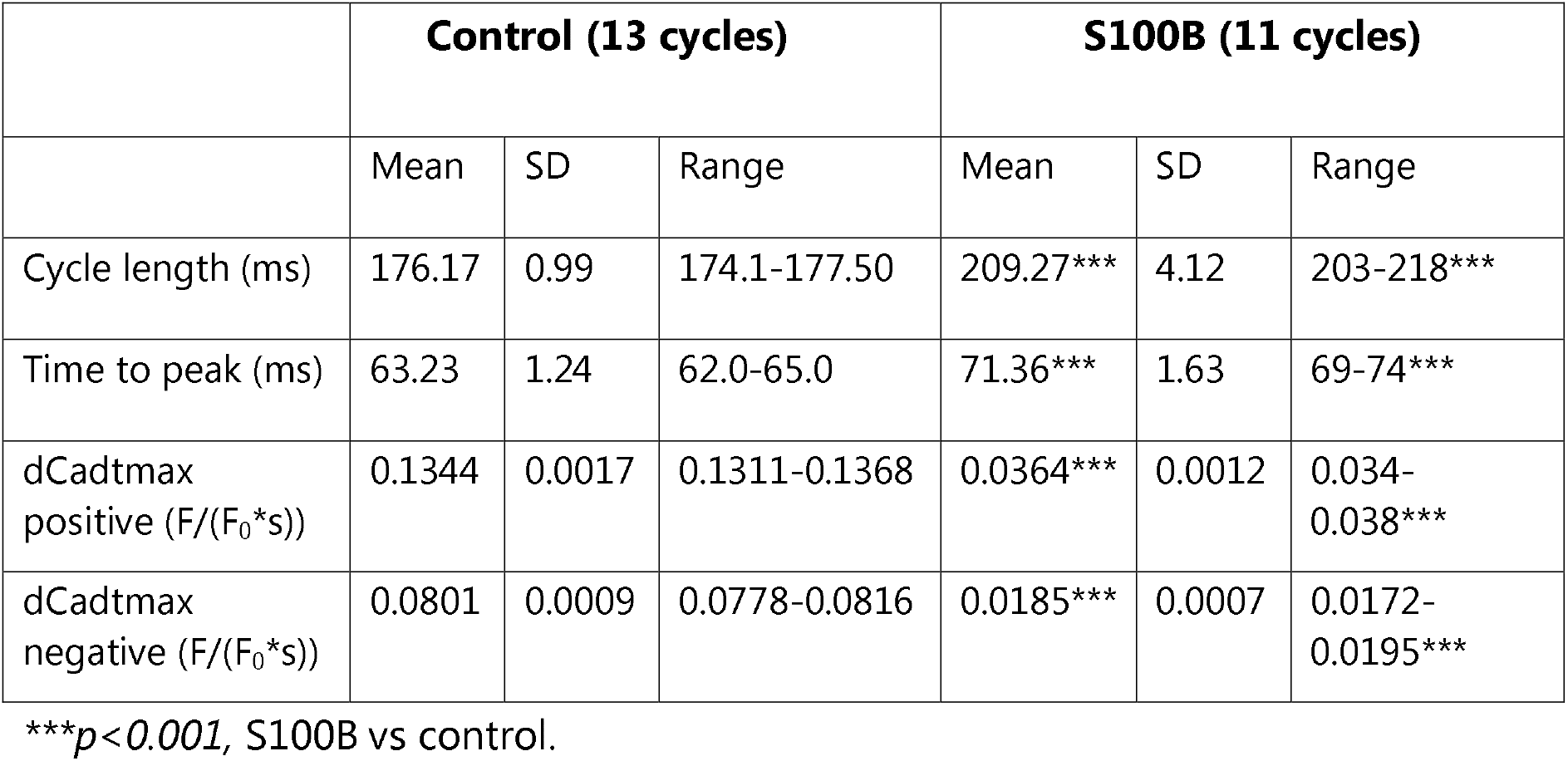
Experiment 2: Ca^2+^ transient parameters in control and during incubation with S100B (mean, SD and range).

|  | Control (13 cycles) |  |  | S100B (11 cycles) |  |  |
| --- | --- | --- | --- | --- | --- | --- |
|  | Mean | SD | Range | Mean | SD | Range |
| Cycle length (ms) | 176.17 | 0.99 | 174.1-177.50 | 209.27*** | 4.12 | 203-218*** |
| Time to peak (ms) | 63.23 | 1.24 | 62.0-65.0 | 71.36*** | 1.63 | 69-74*** |
| dCadtmax positive (F/(F<sub>0</sub>*s)) | 0.1344 | 0.0017 | 0.1311-0.1368 | 0.0364*** | 0.0012 | 0.034-0.038*** |
| dCadtmax negative (F/(F<sub>0</sub>*s)) | 0.0801 | 0.0009 | 0.0778-0.0816 | 0.0185*** | 0.0007 | 0.0172-0.0195*** |

\*\*\* $p < 0.001$ , S100B vs control.

**Supplemental Table 3.** Experiment 3: Ca^2+^ transient parameters in control and during incubation with S100B (mean, SD and range).

|  | Control (12 cycles) |  |  | S100B (8 cycles) |  |  |
| --- | --- | --- | --- | --- | --- | --- |
|  | Mean | SD | Range | Mean | SD | Range |
| Cycle length (ms) | 208.54 | 4.06 | 198.5-213.6 | 292.95*** | 3.32 | 289-297*** |
| Time to peak (ms) | 71.83 | 3.21 | 66.0-77.0 | 82.75*** | 8.81 | 67-97*** |
| dCadtmax positive (F/(F<sub>0</sub>*s)) | 0.0083 | 0.0005 | 0.0078-0.0092 | 0.0067*** | 0.0007 | 0.0054-0.0075*** |
| dCadtmax negative (F/(F<sub>0</sub>*s)) | 0.0042 | 0.0005 | 0.0032-0.0049 | 0.0034*** | 0.0003 | 0.0031-0.0040*** |

\*\*\* $p < 0.001$ , S100B vs control.

**Supplemental Table 4.**
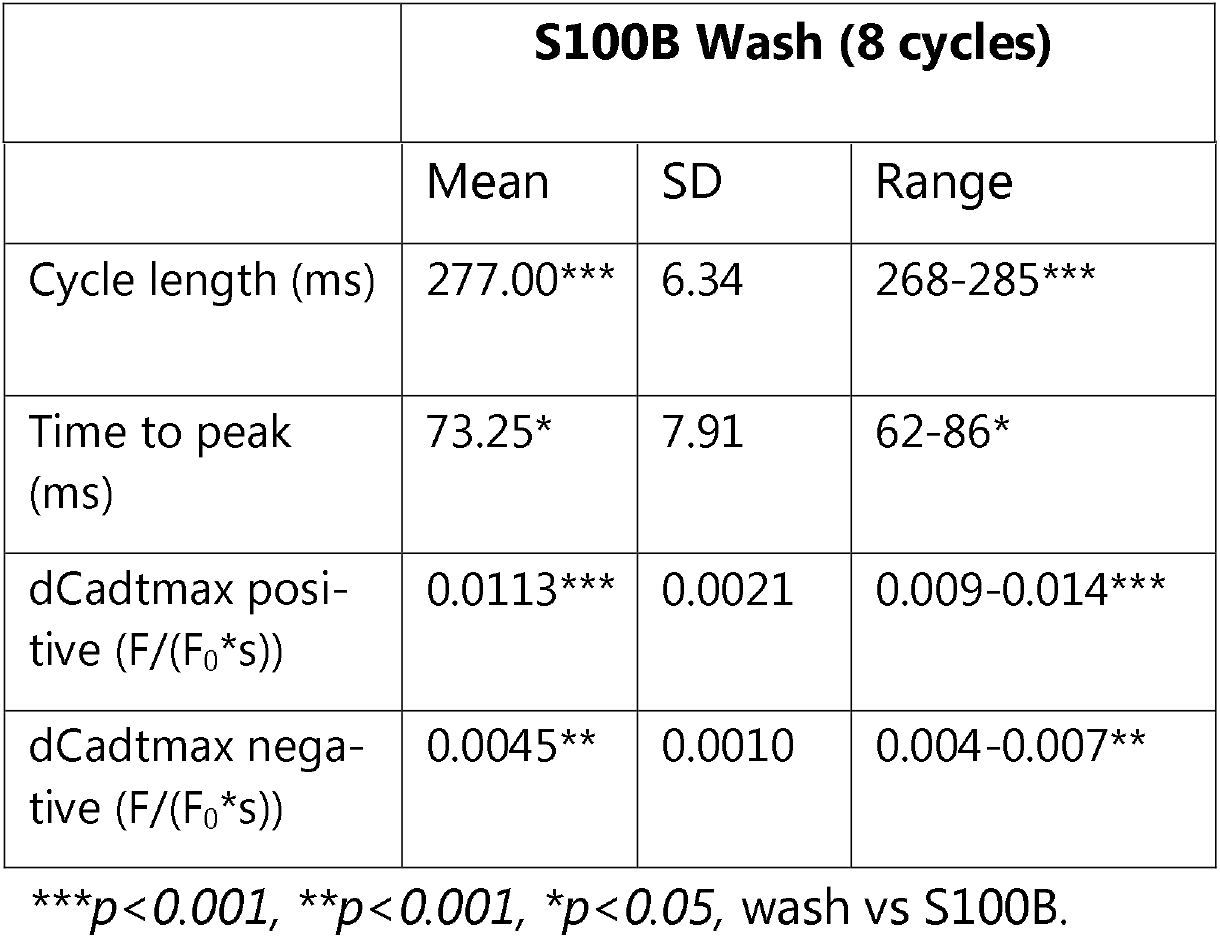
Experiment 3: S100B washout.

## Glossary of Sinoatrial Node Cytoarchitecture

Neuronal fiber: A long projection from a neuronal cell body, an axon
Neuropil: An area between neuronal and glial somata, densely packed with axons, dendrites, and glial processes.
Nerve: A bundle of neuronal fibers, or axons.
Neurite: A single dendritic projection from a neuronal cell body.
Neuronal Plexus: A nerve network. e.g., cholinergic and adrenergic nerves.
Ganglion: A bundle of neuronal somata, often associated with nerve fibers. e.g., the ICG
Varicosity: Postganglionic terminals of autonomic neuronal fibers, site of neurotransmitter release to target organs.
Mesh/Meshwork: A non-hierarchical assembly of heterogeneously distributed and interconnected units. e.g., the HCN4+/CX43- meshwork
Net/Network: A repeating pattern of connections among regularly distributed units. e.g., the CX43+/HCN4- network which locally approaches the HCN4+ meshwork
Web: e.g., the web of peripheral glial cells (PGCs) in the SAN

## Notes

### Competing Interest Statement

The authors have declared no competing interest.

